# Targeting NAT10 suppresses T_FH_ responses and alleviates allergic asthma via an ac4C-TOX2 axis

**DOI:** 10.64898/2026.07.21.739954

**Authors:** Zhou-Li Cheng, Zihao Lin, Yilin Ren, You Zhou, Ying Yang, Zhicheng Yang, Ke Wang, Xushuo Li, Yuhui Zheng, Chuanyu Liu, Xiangdong Wang, Duojiao Wu

**Affiliations:** Jinshan Hospital Center for Tumor Diagnosis & Therapy, Jinshan Hospital, Fudan University, Shanghai, China; Shenzhen Proof-of-Concept Center of Digital Cytopathology, BGI Research, Shenzhen 518083, China; Shanghai Key Laboratory of Lung Inflammation and Injury, Zhongshan Hospital, Fudan University, Shanghai, China

## Abstract

Chemical modifications of RNA, such as N4-acetylcytidine (ac⁴C), fine-tune gene expression, but their roles in cell fate determination within the immune system remain poorly understood. Here, we identify the RNA cytidine acetyltransferase NAT10 as a pivotal regulator of T follicular helper (T_FH_) cell differentiation and the germinal centre response. T cell-specific ablation of Nat10 in mice severely impaired T_FH_ cell development, germinal centre formation, and antibody production following viral infection. Integrated epitranscriptomic and transcriptional profiling revealed that NAT10 deposits ac⁴C modifications on a cohort of mRNAs critical for immune function. We pinpointed the transcription factor TOX2 as a key downstream target, showing that ac⁴C modification within its mRNA coding sequence enhances both transcript stability and translation efficiency, thereby promoting TOX2 protein expression. Enforced TOX2 expression fully rescued the T_FH_ differentiation defect in NAT10-deficient cells. This NAT10-ac⁴C-TOX2 axis is conserved in humans, where its activity correlates with the magnitude of vaccine-elicited T_FH_ responses. Furthermore, we identified R428 as a potent NAT10 inhibitor and demonstrated that its administration could suppress pathogenic T_FH_ responses and alleviate symptoms in a mouse model of allergic asthma. Our work elucidates a central epitranscriptomic mechanism controlling T helper cell fate and humoral immunity, and nominates NAT10 as a potential therapeutic target for T_FH_-driven diseases.

## Introduction

T follicular helper (T_FH_) cells represent a specialized subset of CD4 helper T cells localized within the B cell follicles of secondary lymphoid organs, playing a central role in orchestrating humoral immune responses^[1–3]^. By secreting cytokines such as IL-21 and expressing co-stimulatory molecules including CD40L and ICOS, T_FH_ cells facilitate the selection, affinity maturation, class switching, and differentiation of germinal center (GC) B cells into memory B cells and long-lived plasma cells^[4, 5]^. Functional T_FH_ cells are essential for effective vaccine responses and the establishment of durable immunological memory. Abnormalities in T_FH_ cell function or expansion have been implicated in a wide range of human diseases^[3]^. In autoimmune diseases such as systemic lupus erythematosus and rheumatoid arthritis, hyperactive T_FH_ cells can promote the generation of high-affinity autoreactive antibodies^[6]^. In contrast, dysfunction or dysregulation of T_FH_ cells during chronic infections, including HIV and HBV, can impair viral clearance^[7]^. Therefore, elucidating the regulatory mechanisms underlying T_FH_ cell differentiation is critical for understanding immune homeostasis and the pathogenesis of diverse immunological disorders.

The differentiation of T_FH_ cells is a dynamic and tightly regulated process that requires the integration of multiple signals. This developmental pathway typically proceeds through three sequential stages: initial activation by dendritic cells, migration to the T-B cell border with subsequent interactions with B cells, and terminal differentiation within the germinal center. A range of cytokines and co-stimulatory signals, such as ICOS-ICOSL, CD40-CD40L, IL-6, IL-21, and IL-12, are pivotal in orchestrating this process^[8–10]^. Transcriptional control is central to T_FH_ lineage specification, with Bcl6 functioning as the master regulator^[11]^. Additional transcription factors including TCF1, LEF1, BATF, and others collaboratively drive the expression of T_FH_ signature molecules such as CXCR5 and PD-1^[12–14]^. Meanwhile, repressive factors such as Blimp1, FOXO1, and KLF2 serve as negative regulators, establishing a delicate balance of positive and negative signals^[15]^. Moreover, T_FH_ cells exhibit considerable plasticity, giving rise to functional subsets (e.g., T_FH1_, T_FH2_, T_FH17_) under distinct disease or infection contexts, further underscoring the complexity of their differentiation program.

Despite extensive insights into transcriptional and signaling mechanisms governing T_FH_ cell development, important questions remain unresolved, particularly regarding post-transcriptional regulation. Notably, the potential involvement of RNA modifications in modulating T_FH_ differentiation is largely unexplored. In recent years, epitranscriptomic modifications such as N6-methyladenosine (m⁶A) and N4-acetylcytidine (ac⁴C) have emerged as critical regulators of immune cell development, homeostasis, and inflammatory responses^[16–18]^. Among these, ac⁴C is a conserved RNA modification that enhances mRNA stability and translational efficiency^[19]^. However, whether ac⁴C modification contributes to T_FH_ cell fate decisions and the molecular mechanisms involved remain largely unknown. As such, this represents a promising new frontier at the intersection of RNA biology and T cell immunology, with the potential to reveal novel regulatory paradigms of immune cell differentiation.

In this study, we aimed to investigate whether and how Nat10-mediated ac⁴C modification regulates T_FH_ cell differentiation and antiviral function. Using a conditional knockout mouse model with T cell-specific deletion of Nat10, we observed a marked reduction in T_FH_ cell frequency and impaired germinal center B cell responses following viral infection, suggesting a pivotal role for Nat10 in T_FH_ function. We identified TOX2 mRNA as a direct target of ac⁴C modification and demonstrated that Nat10 deficiency reduces TOX2 ac⁴C levels, leading to decreased mRNA stability and impaired expression. Restoration of TOX2 expression rescued T_FH_ cell differentiation and function in Nat10-deficient mice. Collectively, our findings uncover a novel Nat10–ac⁴C–TOX2 regulatory axis critical for T_FH_ cell fate determination and provide a conceptual framework for targeting RNA modifications to modulate T_FH_ -mediated immunity.

## Results

### Conditional ablation of NAT10 in T cells impairs germinal centre responses

Follicular helper T (T_FH_) cells are distinguished from other CD4^+^ T helper subsets by their role in orchestrating germinal center responses and promoting B cell immunity. In mice immunized intraperitoneally with ovalbumin and alum, T_FH_ cell development correlates positively with IgE and IgG1 production^[20]^. However, whether T_FH_ cells regulate IgE responses in allergic airway inflammation induced by airborne allergens remains poorly understood. To address this, we performed single-cell RNA sequencing (scRNA-seq) on T helper cells isolated from a murine model of allergic airway disease. Our analysis revealed that CD4^+^ T cells could be classified into six distinct subsets. As expected, T_H_2 cells were prevalent in the asthma model, constituting approximately 20.7% of the CD4^+^ T cell population. T_FH_ cells accounted for 10.9% of total CD4^+^ T cells (Figure 1A–B). T_FH_ cells are known to play a critical role in the pathogenesis of asthma, particularly in the generation of allergen-specific IgE antibodies^[21]^. Given our previous finding that NAT10 is essential for CD8^+^ T cell proliferation and antiviral function^[22]^, we sought to investigate its potential role in T helper cell subsets. We therefore examined NAT10 expression across distinct CD4^+^ T cell populations and found that NAT10 was relatively highly expressed in T_FH_ cells compared to other subsets (Figure 1C). To comprehensively investigate the role of *Nat10* in T_FH_ cell differentiation, we generated conditional *Nat10* knockout mice by crossing *Nat10*-floxed mice with *Cd4*-Cre mice transgenic mice. This cross resulted in a T cell-specific deletion of *Nat10* (*Nat10*^fl/fl^*Cd4*^Cre^ mice), enabling the dissection of Nat10’s contribution to T_FH_ cell biology. The efficiency of *Nat10* deletion in splenic CD4^+^ T cells was confirmed through western blot analysis, where a marked reduction in NAT10 protein levels was observed in the *Nat10*^fl/fl^*Cd4*^Cre^ mice (Figure S1A), validating the deletion strategy. To evaluate the functional consequences of *Nat10* loss in vivo, we subjected both *Nat10*^fl/fl^*Cd4*^Cre^ mice and their wild-type littermates (Ctrl) to LCMV-Armstrong infection, a well-established model of acute viral infection that elicits robust T_FH_ and plasma cell responses^[23–25]^. T_FH_ cell differentiation and the formation of plasma cells were monitored at days 3 and 8 post-infection, to capture both the early and late stages of immune response. As anticipated, splenomegaly, a hallmark of viral infection, was observed in control group. However, the spleens of *Nat10*^fl/fl^*Cd4*^Cre^ mice were significantly smaller than those of the control group (Figure 1D), and this was accompanied by a notable reduction in total spleen cell counts at days 3 and day 8 post-infection (Figure 1E), suggesting impaired immune cell proliferation in the spleen following the loss of NAT10. The most striking finding was a marked reduction in the frequency and total number of CD44^+^CXCR5^+^ T_FH_ cells in the spleens of *Nat10*^fl/fl^*Cd4*^Cre^ mice at day 8 post-infection, relative to controls (Figures 1F-G). Additionally, a significant decrease in the total number of CD44^+^CXCR5^+^ T_FH_ cells was observed in the spleens of *Nat10*^fl/fl^*Cd4*^Cre^ mice as early as day 3 post-infection (Figures 1F-G). However, at day 8 post-infection, the frequency of CD44^+^CXCR5^-^ T_H_1 cells in the spleens of *Nat10*^fl/fl^*Cd4*^Cre^ mice is significantly higher compared to the controls, while the number in *Nat10*^fl/fl^*Cd4*^Cre^ mice is significantly lower (Figure S1B). This observation was further corroborated by a detailed analysis of germinal center (GC) T_FH_ cells, which play a pivotal role in providing help to B cells for antibody production. We observed that the absence of NAT10 resulted in a profound reduction in both the percentage and the total number of GC T_FH_ cells, as identified by the canonical markers PD-1^hi^CXCR5^+^, ICOS^hi^CXCR5^+^, or Bcl-6^hi^CXCR5^+^ (Figures 1H-I, S1C-D). Notably, the expression levels of these critical T_FH_-associated markers—CXCR5, PD-1, ICOS, and Bcl-6—were significantly diminished on T_FH_ cells from *Nat10*^fl/fl^*Cd4*^Cre^ mice compared to controls, further underscoring the essential role of NAT10 in the differentiation and functional regulation of T_FH_ cells (Figure S1E). No significant differences were observed between *Nat10*^fl/fl^ and *Nat10*^fl/fl^*Cd4*^Cre^ mice at the double-negative, double-positive, and mature single-positive stages, respectively (Figures S2A-B), indicating that early T cell development in the thymus remains unaffected. Furthermore, we analyzed the changes in cell proportions in the spleen and found that the proportion of CD4^+^ and CD8^+^ T cells was significantly reduced by approximately threefold in Nat10-deficient mice, consistent with our previous findings (Figures S2C-D). We also assessed the distribution of different CD4^+^ T cell subsets. Prior to LCMV infection, *Nat10* knockout resulted in a significant increase in the proportion of T_H_1 cells, indicative of an inflammatory state. Correspondingly, the proportion of T_H_2 cells also significantly increased, in line with our earlier observations^[26, 27]^ (Figures S2E-F). Post-LCMV infection, *Nat10* knockout led to a marked decrease in the proportion of T_H_1 cells, while the proportion of T_H_2 cells remained unchanged (Figures S2E-F). The proportion of T_H_17 cells was low in the spleen T cell population, and neither *Nat10* knockout nor LCMV infection had a notable impact on this subset (Figures S2G). The proportion of Treg cells decreased following LCMV infection, but no significant change was observed in *Nat10*-deficient mice (Figures S2H). We also performed *in vitro* T_FH_ differentiation assays to assess the impact of Nat10 on T_FH_ differentiation. CD4^+^ naïve T cells were sorted from the spleens of *Nat10*^fl/fl^ and *Nat10*^fl/fl^*Cd4*^Cre^ mice and induced to differentiate into T_FH_ cells. Consistent with the *in vivo* results, we observed that Nat10 knockout significantly impaired T_FH_ cell differentiation in the *in vitro* differentiation assay (Figures S3A-D). Additionally, we evaluated the proliferative capacity of T_FH_ cells. CD4^+^ naïve T cells were labeled with carboxyfluorescein diacetate succinimidyl ester (CFSE) and cultured under differentiation conditions for 72 hours. T_FH_ cell proliferation was measured, and no significant difference was observed in Nat10-deficient T_FH_ cells, suggesting that the reduced frequency of T_FH_ cells observed *in vitro* is due to a diminished differentiation capacity rather than a decrease in cell proliferation(Figures S3E-F).

**Figure 1.**
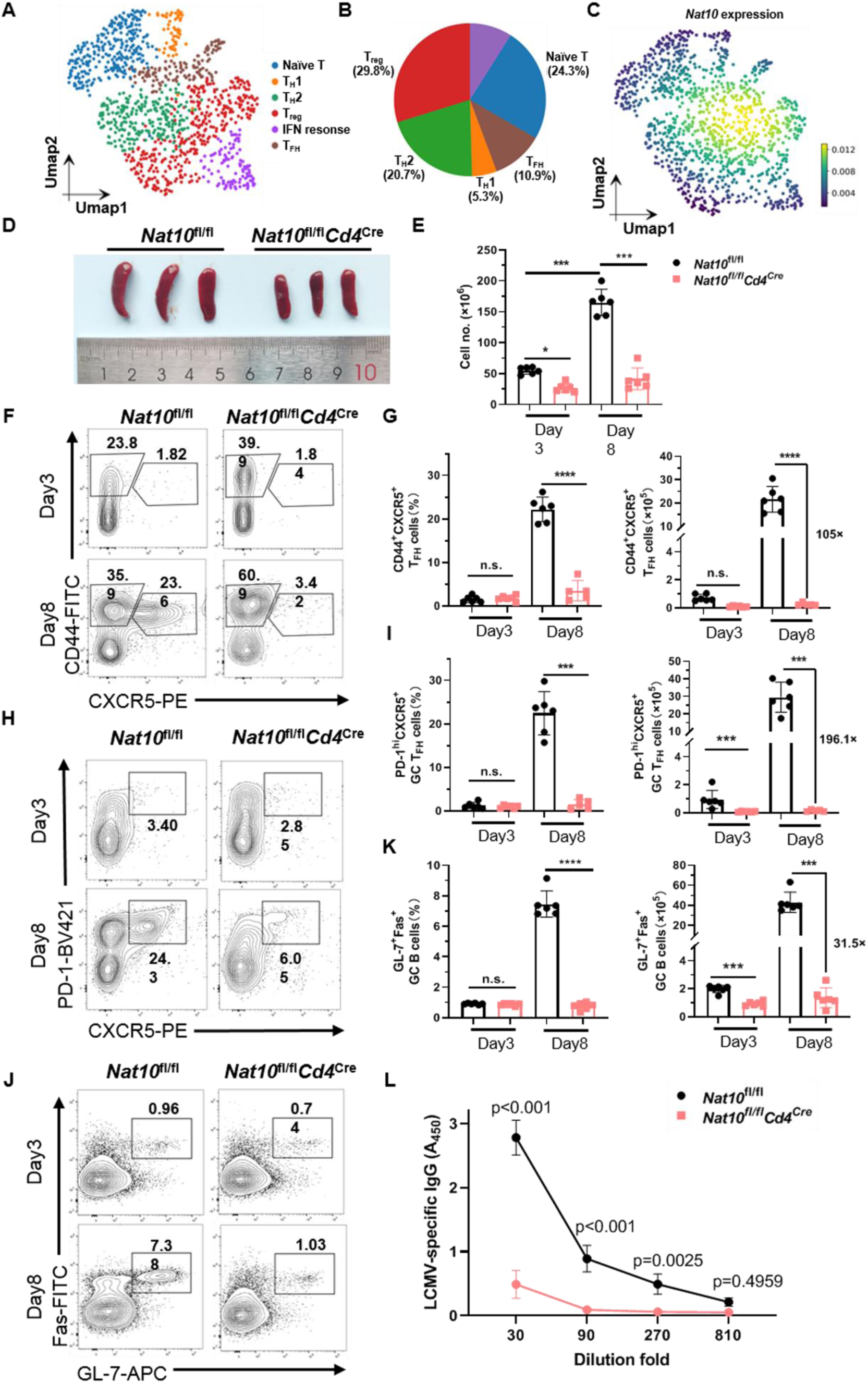
Conditional ablation of Nat10 impairs T_FH_ differentiation and germinal centre responses. **A**, UMAP visualization of all CD4^+^ T cells isolated from the lungs and lymph nodes of mice with allergic airway disease (GSE131935). **B,** Pie chart showing the distribution of distinct T helper cell subsets. **C,** Nat10 expression across different T helper cell populations. **D**, Representative spleen images from LCMV-Armstrong-infected *Nat10*^fl/fl^ and *Nat10*^fl/fl^*Cd4*^Cre^ mice at day 8 post-infection. **E,** Total splenocyte counts from infected mice at days 3 and 8 p.i.. **F-G**, Representative flow cytometry plots of CD44^+^CXCR5^+^ T_FH_ and CD44^+^CXCR5^-^ T_H_1 cells, gated on splenic CD4^+^ T cells, from *Nat10*^fl/fl^ and *Nat10*^fl/fl^*Cd4*^Cre^ mice at days 3 and day 8 post-infection (**F**). Frequency and absolute numbers of indicated cell subsets at days 3 and day 8 post-infection (n = 6 per group) (**G**). **H-I**, Representative flow cytometry plots of PD-1^hi^CXCR5^+^ GC T_FH_ cells, gated on splenic CD4^+^ T cells, from *Nat10*^fl/fl^ and *Nat10*^fl/fl^*Cd4*^Cre^ mice at days 3 and day 8 post-infection (**H**). Frequency and absolute numbers of indicated cell subsets at days 3 and day 8 post-infection (n = 6 per group) (**I**). **J-K**, Representative flow cytometry plots of GL-7^+^Fas^+^ GC B cells from *Nat10*^fl/fl^ and *Nat10*^fl/fl^*Cd4*^Cre^ mice at days 3 and day 8 post-infection (**J**). Frequency and absolute numbers of indicated cell subsets at days 3 and day 8 post-infection (n = 6 per group) (**K**). **L,** Serum LCMV-specific IgG titres from *Nat10*^fl/fl^ and *Nat10*^fl/fl^*Cd4*^Cre^ mice at days 8 post-infection were measured by ELISA (n = 4 per group). P values are calculated using two-tailed Student’s t-test for paired comparisons or one-way ANOVA for multiple comparisons. *denotes p < 0.05, ***denotes p < 0.001, and ****denotes p < 0.0001 for the indicated comparison. n.s. = not significant.

Given the essential role of T_FH_ cells in facilitating B cell activation and germinal center (GC) formation—key processes underlying humoral immunity and the establishment of long-term immune memory—we next sought to investigate the consequences of NAT10 deficiency on GC B cell responses. In *Nat10*^fl/fl^*Cd4*^Cre^ mice, we observed a striking reduction in both the proportion and absolute number of GL-7^+^Fas^+^ GC B cells, key indicators of GC activation (Figure 1J-K). Additionally, a significant decrease in the total number of GL-7^+^Fas^+^ GC B cells was observed in the spleens of *Nat10*^fl/fl^*Cd4*^Cre^ mice as early as day 3 post-infection (Figure 1 J-K). This reduction was accompanied by a significant decrease in the number of IgD^lo^CD138^+^ plasma cells (Figure S1F), which are crucial for the generation of antibody-secreting cells. These observations suggest that the loss of NAT10 impairs the formation and maintenance of functional GC responses, thereby compromising B cell maturation. To further assess the impact of this impaired T_FH_ and GC cell response on humoral immunity, we measured the concentration of LCMV-specific IgG in the serum of infected mice. Notably, *Nat10*^fl/fl^*Cd4*^Cre^ mice exhibited significantly reduced levels of LCMV-specific IgG at day 8 post-infection, compared to their wild-type littermates (Figure 1L). This reduction in antibody production underscores the pivotal role of NAT10 in the orchestration of a robust humoral immune response, highlighting its importance in the regulation of T_FH_ cell differentiation and GC B cell activity. The absence of NAT10 leads to defective humoral immunity, impairing the ability to mount a strong and sustained immune response following viral infection.

Together, these data strongly suggest that NAT10 is a critical regulator of T_FH_ cell differentiation, playing an indispensable role in the generation and maintenance of T_FH_ cells during immune responses to viral infection.

### NAT10 orchestrates the transcriptional profiles of T_FH_ cells

We next sought to investigate the impact of NAT10 deficiency on the transcriptional profiles of T_FH_ cells. CD4^+^ T cells typically differentiate into T_H_1 or T_FH_ cells upon acute viral infection, and since NAT10 deficiency impairs CXCR5 expression, we employed CD44 and SLAM as markers to distinguish between T_H_1 and T_FH_ cells in activated CD4^+^ T cells. Notably, SLAM is expressed at low levels on T_FH_ cells, making it an ideal marker for their identification. We isolated CD44^+^SLAM^hi^ T_H_1 cells and CD44^+^SLAM^lo^ T_FH_ cells from *Nat10*^fl/fl^*Cd4*^Cre^ mice and their corresponding Ctrl littermates at day 8 post-viral infection, and performed RNA-seq analysis on these populations (Figure S4A).

Principal component analysis (PCA) and hierarchical clustering revealed substantial differences in the transcriptional profiles of T_H_1 and T_FH_ cells between *Nat10*^fl/fl^*Cd4*^Cre^ mice and their Ctrl littermates, highlighting the pivotal role of NAT10 in shaping the transcriptome of T cells (Figure 2A). This result indicates that the absence of NAT10 leads to significant transcriptional changes in both T_H_1 and T_FH_ cells. Through gene expression clustering analysis of the RNA-seq data, we identified four distinct gene clusters in response to viral infection (Figure 2B and Table S1). Cluster IV genes were upregulated in both T_H_1 and T_FH_ cells following NAT10 deletion and included several genes involved in the inflammatory response, such as Cxcl10, Cxcl9, and Ccr7. This finding is consistent with our previous reports indicating that NAT10 deficiency promotes an upregulation of inflammatory responses in mice^[22]^. In contrast, genes within Cluster I were downregulated in NAT10-null T cells and included several adaptive immune response genes such as Tox2, Pdcd1, Tcf7, and Icos, all of which play key roles in T_FH_ cell differentiation. Upon examining the gene expression profiles in more detail, we found 1094 upregulated genes and 480 downregulated genes in NAT10-deficient T_FH_ cells (Figure 2C and Table S2), and 1690 upregulated genes and 598 downregulated genes in NAT10-deficient T_H_1 cells (Figure S4B), with ≥2-fold expression changes and an adjusted P-value <0.01. Notably, the downregulated genes in T_FH_ cells were significantly enriched in the T cell receptor signaling pathway and the adaptive immune response(Figure 2D), whereas the upregulated genes in T_FH_ cells were primarily associated with immune system processes and inflammatory responses (Figure S4C). In T_H_1 cells, downregulated genes were enriched in cell division and adaptive immune responses, while upregulated genes were more prominently involved in immune system processes and innate immunity (Figure S4D-E). Next, we performed Gene Set Enrichment Analysis (GSEA) using both T_FH_ cell and GC T_FH_ cell signature gene sets^[12]^. Both gene sets were significantly enriched in wild-type T_FH_ cells but showed no enrichment in NAT10-deficient T_FH_ cells (Figure 2E and S4F). Similarly, GSEA of T_H_1^[28]^, T_H_2^[29]^, T_H_17^[30]^, and Treg^[29]^ signature gene sets revealed that both T_H_1 and T_H_17 gene sets were enriched in wild-type T_FH_ cells, but not in their NAT10-deficient counterparts (Figure S4G). A parallel analysis of T_H_1 cells showed similar results, with T_H_1 and T_H_17 gene sets enriched in wild-type T_H_1 cells, but not in NAT10-deficient T_H_1 cells (Figure S4H). Collectively, these findings underscore that the absence of NAT10 disrupts the transcriptional programs of both T_FH_ and T_H_1 cells, resulting in disordered gene expression profiles in both subsets.

**Figure 2.**
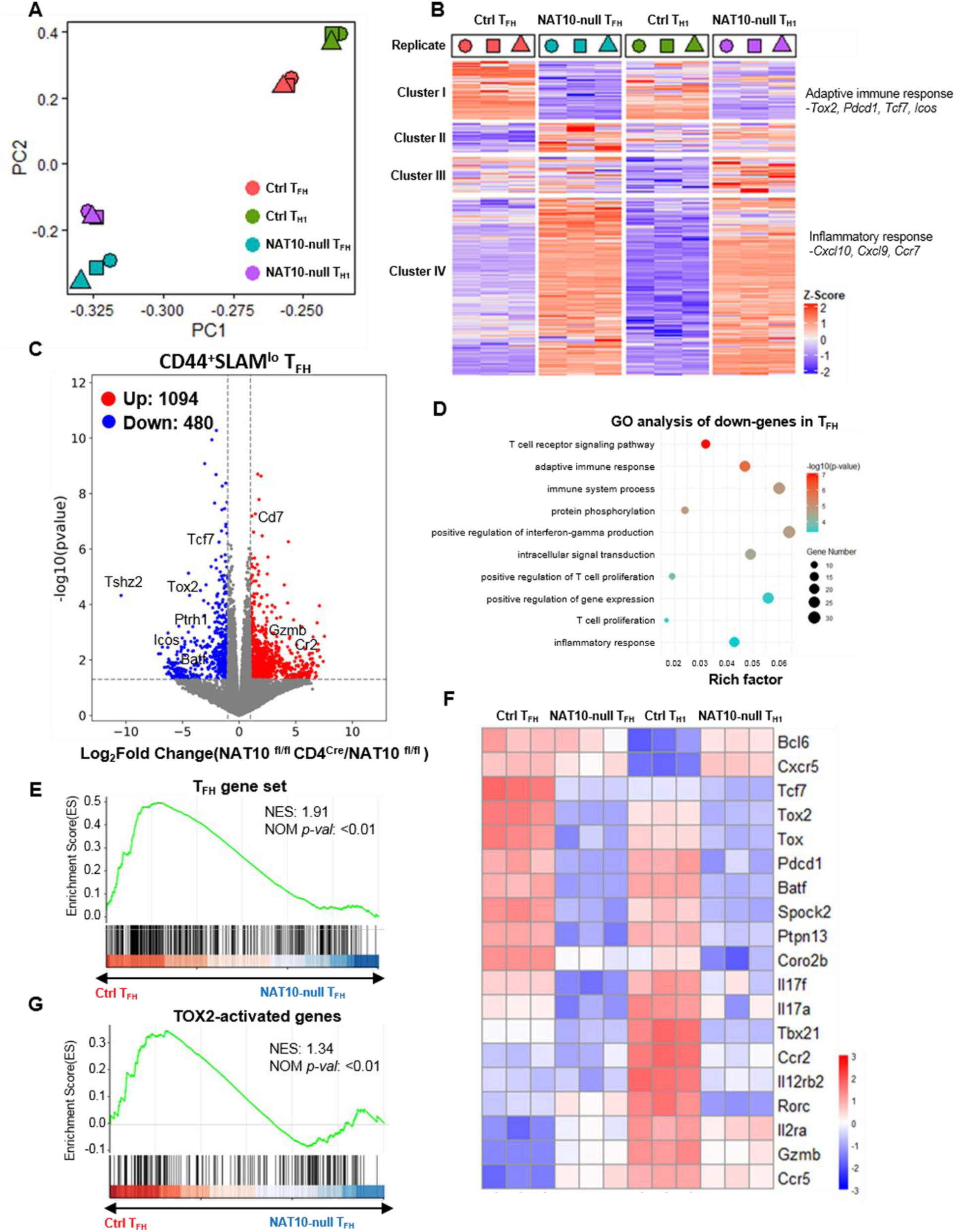
Altered transcriptional profiles in NAT10-deficient T_FH_ cells. **A**, Principal component analysis (PCA) of RNA-seq data from sorted CD44^+^SLAM^hi^ T_H_1 and CD44^+^SLAM^lo^ T_FH_ cells at day 8 p.i. Each group contains three biological replicates. **B,** Heatmap and clustering analysis of genes differentially expressed among T_H_1 cells and T_FH_ cells from *Nat10*^fl/fl^ and *Nat10*^fl/fl^*Cd4*^Cre^ mice at day 8 post-viral infection. The circle, square and triangle symbols stand for three biological replicates. **C**, Volcano plot showing differentially expressed genes between *Nat10*^fl/fl^ and *Nat10*^fl/fl^*Cd4*^Cre^ T_FH_ cells at day 8 post-viral infection. All differentially expressed genes with P<0.05 and fold change>2 are highlighted in red (up-regulation) and blue (down-regulation). **D**, Top GO enrichment analysis of the down-regulated genes in *Nat10*^fl/fl^*Cd4*^Cre^ T_FH_ cells compared with *Nat10*^fl/fl^ T_FH_ cells. **E**, Gene set enrichment analysis (GSEA) of TFH gene set in *Nat10*^fl/fl^*Cd4*^Cre^ T_FH_ cells relative to expression in *Nat10*^fl/fl^ T_FH_ cells. **F**, Heatmaps representation of T_FH_ associated genes in the *Nat10*^fl/fl^ and *Nat10*^fl/fl^*Cd4*^Cre^ T cells are shown, respectively. **G,** GSEA analysis of “TOX2-activated genes in TFH cells” in *Nat10*^fl/fl^*Cd4*^Cre^ T_FH_ cells relative to expression in *Nat10*^fl/fl^ T_FH_ cells.

To evaluate the validity of using CD44^+^SLAM^lo^ cells as a surrogate for true Tfh cells, we compared the gene expression profiles of CD44^+^SLAM^lo^ cells with those of bona fide T_FH_ cells (CD4^+^CD44^+^CXCR5^+^BCL6^+^) from the previous study ^[31]^. Our analysis revealed that CD44^+^SLAM^lo^ cells exhibit a gene expression profile highly similar to that of true T_FH_ cells. In contrast, CD44^+^SLAM^hi^ cells displayed an expression profile more akin to that of T_H_1 cells (Figure S5A). To further validate these findings, we assessed the expression of key T_FH_-associated genes and observed that *Tcf7*, *Pdcd1*, and *Icos* were significantly downregulated in NAT10-null T_FH_ cells compared to wild-type controls, while *Bcl6* and *Cxcr5* did not show significant changes (Figure 2F). Interestingly, following Nat10 knockout, protein levels of BCL6 and CXCR5 were significantly reduced. We hypothesize that Nat10 may affect the translation efficiency of *Bcl6* and *Cxcr5*. To explore this, we performed polyribosome real-time PCR experiments to quantify ribosome occupancy on *Bcl6* and *Cxcr5* mRNAs. Notably, we observed a marked reduction in ribosome accumulation on *Bcl6* and *Cxcr5* mRNAs, but not on control transcripts, upon Nat10 depletion (Figure S5B-C). Additionally, *Tox2*, a gene known to be involved in T_FH_ cell differentiation and regulate *Tcf7*, *Pdcd1*, and *Icos* expression^[32]^, was also expressed at lower levels in NAT10-deficient T_FH_ cells compared to their wild-type counterparts (Figure 2F). Given the lower expression of *Tox2*, we focused our attention on this gene due to its critical role in T_FH_ cell differentiation. To gain further insight, we performed GSEA with a gene set containing TOX2-activated genes in T_FH_ cells^[32]^. Remarkably, the TOX2-associated gene set was enriched in wild-type T_FH_ cells, but not in NAT10-deficient T_FH_ cells (Figure 2G). These results suggest that NAT10 plays a crucial role in regulating the T_FH_ differentiation program by activating genes associated with the T_FH_ lineage. Taken together, our data demonstrate that NAT10 governs the transcriptional network essential for T_FH_ cell differentiation, and its deficiency leads to significant alterations in the gene expression profile of both T_FH_ and T_H_1 cells.

### ac^4^C modifies *Tox2* mRNA to control its stability

To explore how ac^4^C acetylation regulates the transcriptional program of T_FH_ cells, we employed ac^4^C-RIP-seq to map the global ac^4^C landscape of CD4^+^ T cells isolated from either Ctrl or *Nat10*^fl/fl^*Cd4*^Cre^ mice infected with LCMV-Armstrong. Our bioinformatic analysis revealed that ac^4^C peaks were predominantly located within the coding sequences (CDS) and near the start codon of mRNAs (Figure 3A-B), underscoring a pattern of ac^4^C enrichment that aligns with regions critical for translation initiation. Approximately 41.9% of ac^4^C-modified mRNAs contained two or more ac^4^C peaks, highlighting a broad, complex involvement of this modification in mRNA regulation (Figure 3C). From biological triplicates of ac^4^C-RIP-seq, we identified a total of 6405 transcripts that were classified as ac^4^C-modified transcripts (Figure 3D and Table S3), and predicted an octanucleotide motif (CDGCAKC) as the putative ac^4^C acetylation motif (E-value = 1.0e-070) (Figure S6A). Strikingly, when we compared these ac^4^C-modified transcripts with differentially expressed genes, we found that 141 transcripts, including key regulators such as *Tox2* and *Maf*, were potentially modulated by ac^4^C acetylation. Gene Ontology (GO) term analysis further revealed that these transcripts were enriched for biological processes related to positive regulation of transcription and gene expression, highlighting the functional relevance of ac^4^C modification in immune cell regulation (Figure 3E). Upon closer examination of specific mRNAs, we discovered that the CDS of *Tox2* mRNA exhibited highly enriched ac^4^C peaks, whereas other critical T_FH_-related transcripts, such as *Tcf7* and *Icos*, do not exhibit significant changes in ac^4^C modification upon NAT10 deficiency (Figure 3F and S6B-E). These results suggested that *Tox2* is a bona fide target of ac^4^C acetylation. To confirm this finding, we performed RNA immunoprecipitation (RIP) assays, which demonstrated that NAT10 protein directly binds to *Tox2* mRNA (Figure 3G). Additionally, ac^4^C-RIP-qPCR further confirmed that *Tox2* mRNA was indeed tagged by ac^4^C acetylation in CD4^+^ T cells (Figure 3H). Together, these data firmly established *Tox2* as a key ac^4^C-modified transcript. Given that ac^4^C modification regulates gene expression through various mechanisms, including mRNA stability, and translation, we next sought to investigate how ac^4^C modification influences TOX2 expression at the translational level. To do so, we performed polyribosome profiling via real-time PCR and observed a dramatic decrease in ribosome accumulation on *Tox2* mRNA in NAT10-deficient cells, whereas no such reduction was observed for control *Actb* mRNA transcripts (Figure 3I). This suggested that the absence of NAT10 impairs the translation of *Tox2* mRNA. We then examined the stability of *Tox2* mRNA by performing RNA decay assays using actinomycin D (ActD) treatment to block transcription. Interestingly, we observed that *Tox2* mRNA exhibited significantly accelerated degradation in NAT10-deficient cells compared to Ctrl cells, as evidenced by a rapid decline in transcript levels at multiple time points post-treatment (Figure 3J-K). This finding suggested that NAT10 contributes to the stability of *Tox2* mRNA, thereby promoting its expression. To determine whether the reduced *Tox2* mRNA stability and translation affected protein synthesis, we measured the TOX2 protein levels in NAT10-deficient T_FH_ cells. Indeed, we found that TOX2 protein was significantly decreased in NAT10-deficient T_FH_ cells compared to their wild-type counterparts (Figure 3L), confirming that NAT10 plays a critical role in facilitating the synthesis of TOX2 protein. In summary, our findings provide compelling evidence that NAT10 promotes the synthesis of TOX2 protein by enhancing both the stability and translation of *Tox2* mRNA. This regulation is mediated by ac^4^C acetylation, which serves as a key mechanism through which NAT10 modulates T_FH_ cell differentiation and function.

**Figure 3.**
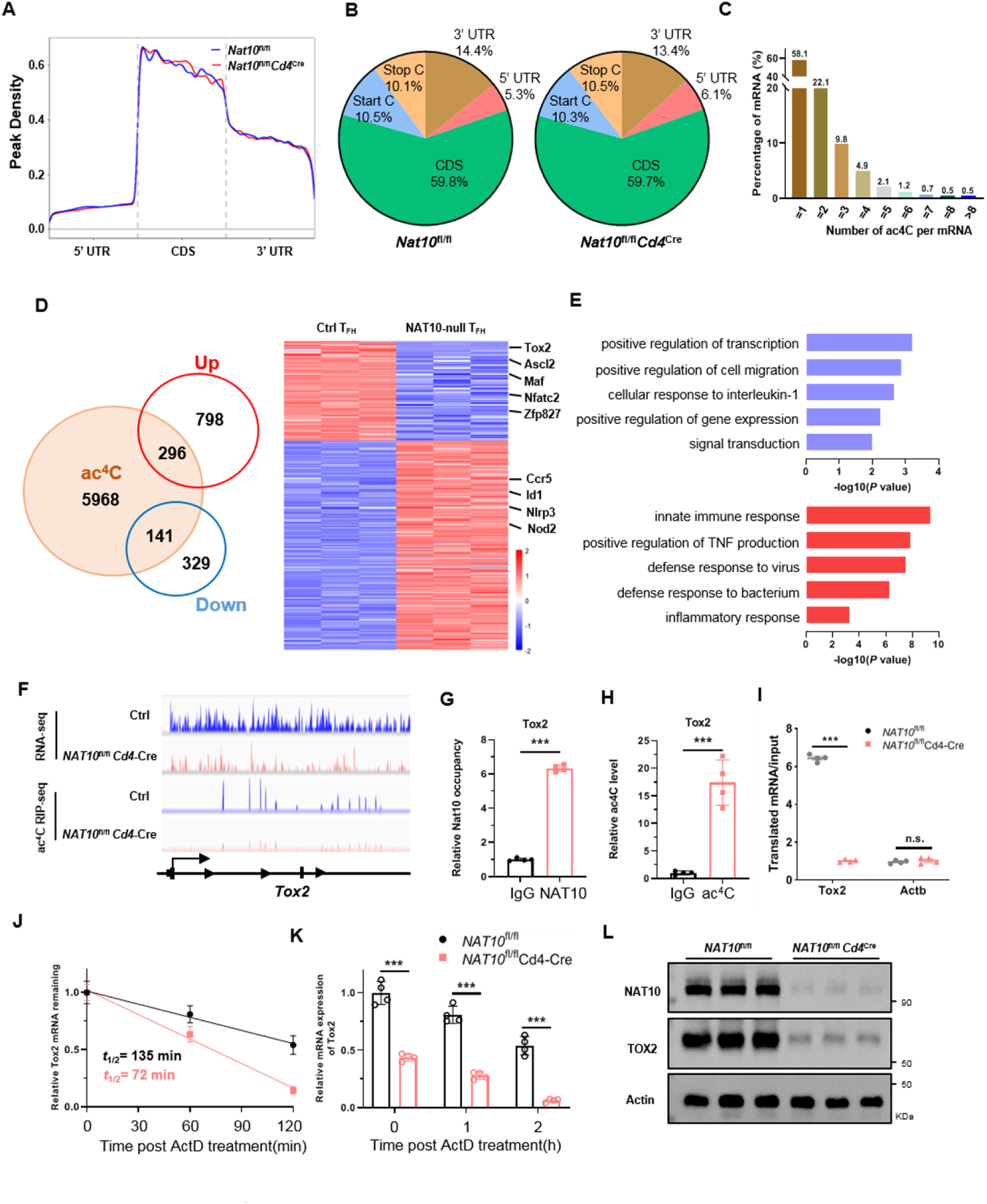
NAT10 and ac^4^C promotes TOX2 protein synthesis by enhancing *Tox2* mRNA stability and translation. **A**, Metagene profiles of ac^4^C site distribution on mRNAs from *Nat10*^fl/fl^ and *Nat10*^fl/fl^*Cd4*^Cre^ T_FH_ cells (UTR, untranslated region; CDS, coding sequence). **B,** Pie chart showing the distribution of ac^4^C sites in five regions of *Nat10*^fl/fl^ and *Nat10*^fl/fl^*Cd4*^Cre^ T_FH_ cells. **C**, Bar chart depicting percentage of mRNAs with different internal ac^4^C abundance. **D**, Venn diagram showing the overlapping ac^4^C-modified transcripts and differentially expressed genes (left). Heatmap of differentially expressed genes with ac^4^C modification in *Nat10*^fl/fl^ and *Nat10*^fl/fl^*Cd4*^Cre^ T_FH_ cells (right). **E**, Top GO terms of the biological process categories enriched in differentially expressed transcripts with ac^4^C peaks (Red: up-regulated genes; blue: down-regulated genes). **F**, Integrative Genomics Viewer (IGV) tracks displaying RNA-seq (top panel) and ac^4^C RIP-seq (bottom panel) reads distribution of *Tox2* gene. **G,** RIP-qPCR analysis showing enrichment of NAT10 on *Tox2* mRNA in T_FH_ cells. The *Tox2* mRNA enrichment is presented as IP/input and normalized to IgG group (n =4 per group). **H,** ac^4^C RIP–qPCR for *Tox2* mRNA in T_FH_ cells (n =4 per group). **I,** Ribosome occupancy of *Tox2* and control *Actb* mRNAs in *Nat10*^fl/fl^ and *Nat10*^fl/fl^*Cd4*^Cre^ T_FH_ cells. (n =4 per group). **J,** Degradation of *Tox2* mRNA in *Nat10*^fl/fl^ and *Nat10*^fl/fl^*Cd4*^Cre^ T_FH_ cells. The half-live of *Tox2* mRNA was detected by quantitative RT-PCR. The remaining mRNAs were normalized to t = 0 (n = 4 per group). **K,** Quantitative RT-PCR analysis of *Tox2* mRNA abundance as in **J**, relative expression was normalized to t = 0 of *Nat10*^fl/fl^ cells (n = 4 per group). **L,** The protein level of NAT10 and TOX2 in *Nat10*^fl/fl^ and *Nat10*^fl/fl^*Cd4*^Cre^ T_FH_ cells, as determined by western-blot. Each group contains three biological replicates. P values are calculated using two-tailed Student’s t-test for paired comparisons or one-way ANOVA for multiple comparisons. ***denotes p < 0.001 for the indicated comparison. n.s. = not significant.

### Forced expression of TOX2 restores defective T_FH_ differentiation in NAT10-deficient mice

To elucidate the functional connection between NAT10 and the stability of *Tox2* transcripts in T_FH_ differentiation, we next sought to investigate the impact of ectopic expression of TOX2 in NAT10-deficient cells. To this end, we engineered a retrovirus vector expressing full-length TOX2 (TOX2-RV-GFP), which was subsequently used to infect activated CD4^+^ T cells. GFP^+^ cells were isolated by fluorescence-activated cell sorting (FACS), and the sorted cells were cultured under T_FH_-like conditions (α-IFNγ+α-IL-4+α-IL-2+IL-6), as previously described^[32]^. Notably, retroviral overexpression of TOX2 led to a substantial upregulation of Bcl6 mRNA, alongside other key T_FH_-associated genes, including Cxcr5, Il6ra, Pdcd1, Il6st, and Tcf7 (Figure 4A), suggesting that TOX2 plays a critical role in promoting T_FH_ differentiation. To assess the in vivo relevance of these findings, we next performed bone marrow chimeric experiments. We transferred TOX2-transduced wild-type (Ctrl) or NAT10-deficient (*Nat10*^fl/fl^*Cd4*^Cre^) CD4^+^ T cells into lethally irradiated recipient mice to generate bone marrow chimeras (Figure 4B). Following infection with LCMV-Armstrong, we analyzed the T_FH_ populations derived from transduced CD4^+^ T cells in recipient mice on day 8 post-infection. As expected, NAT10-deficient CD4⁺ T cells transduced with empty vector (EV) showed impaired differentiation into CD44⁺CXCR5⁺ T_FH_ cells compared with controls (Figure 4C). Importantly, TOX2 overexpression largely rescued this defect, restoring T_FH_ differentiation to levels comparable with controls. These results indicate that TOX2 can compensate for the loss of NAT10 in T_FH_ differentiation. Because our previous work identified Myc as a NAT10 target whose protein synthesis is promoted by NAT10, we asked whether Myc could enhance T_FH_ differentiation. Overexpression of Myc alone in NAT10-deficient CD4⁺ T cells failed to improve T_FH_ differentiation. However, co-expression of Myc and TOX2 significantly enhanced the differentiation efficiency compared with TOX2 alone, suggesting a role in T_FH_ expansion (Figure 4D). Accordingly, germinal center (GC) T_FH_ populations (identified as PD-1^hi^CXCR5^+^, ICOS^hi^CXCR5^+^, or Bcl-6^hi^CXCR5^+^) were restored in TOX2-overexpressing NAT10-deficient cells, with further improvement upon Myc co-expression (Figures 4E–F, 4I–J).

**Figure 4.**
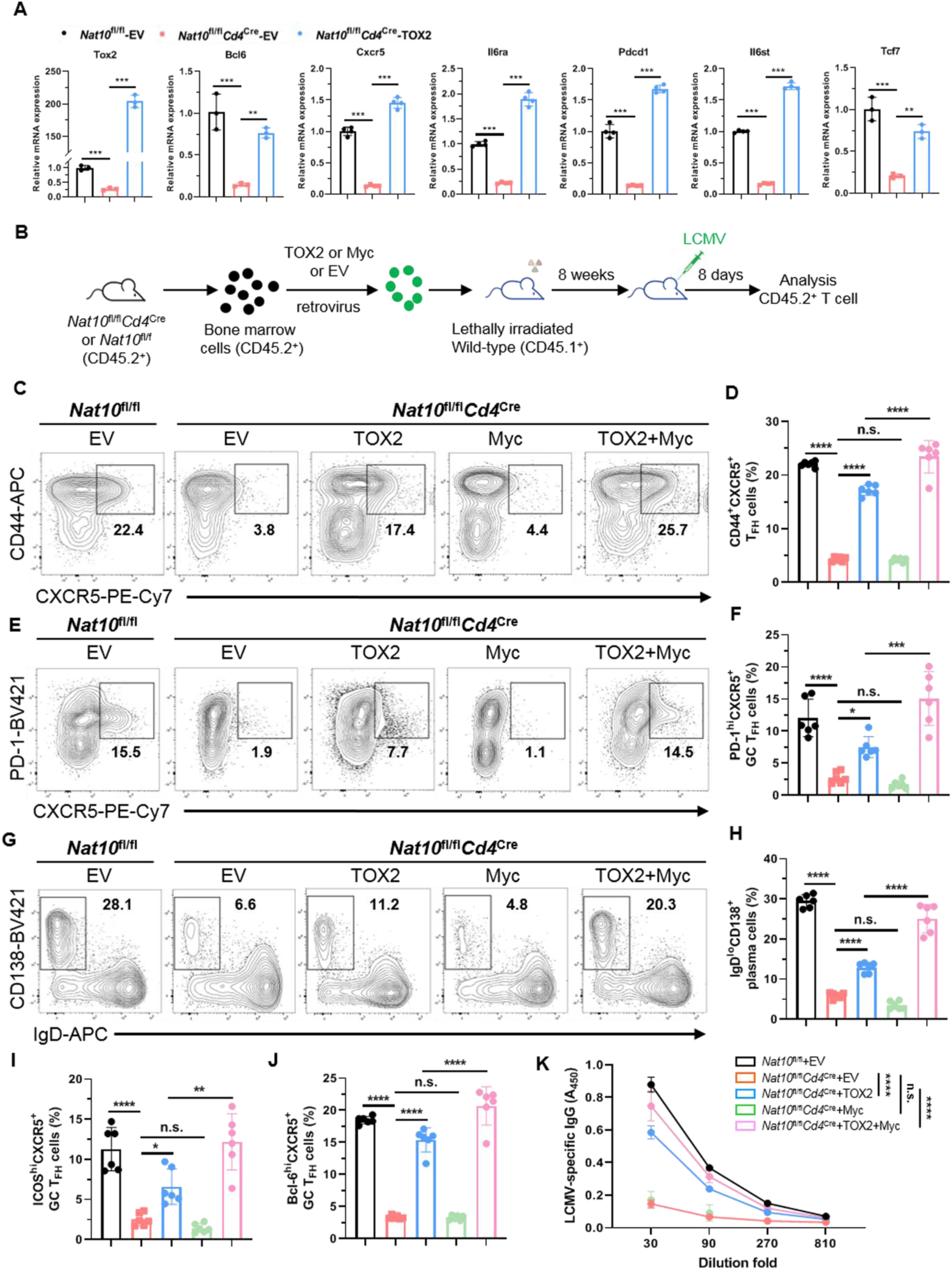
TOX2 overexpression rescues defective T_FH_ differentiation in NAT10-deficient cells. **A**, *Nat10*^fl/fl^ and *Nat10*^fl/fl^*Cd4*^Cre^ T_FH_ cells were infected with empty vector (EV) or TOX-2 retrovirus, and the mRNA expression of Tox2, Bcl6, Cxcr5, Il6ra, Pdcd1, Il6st and Tcf7 genes were determined by RT-qPCR. **B,** Schematic diagram showing construction of bone marrow chimeric mice. CD117^+^ enriched HSCs from 8-week-old *Nat10*^fl/fl^ and *Nat10*^fl/fl^*Cd4*^Cre^ mice were infected with empty vector, Myc or TOX2 retrovirus and then transplanted into irradiated 8-week-old CD45.1^+^ mice. Eighth weeks later, bone marrow chimeric mice were intraperitoneally injected with LCMV Armstrong, and CD45.2^+^ T cells derived from retrovirus-infected HSCs were identified for further analysis. **C-D**, Representative flow cytometry plots of CD44^+^CXCR5^+^ T_FH_ populations gated on adoptive cells identified by CD45.2 staining in each group at 8 days post infection. Frequency of donor-derived T_FH_ cells are shown in **D** (n = 6). **E-F**, Representative flow cytometry plots of PD-1^hi^CXCR5^+^ GC T_FH_ populations gated on adoptive cells identified by CD45.2 staining in each group at 8 days post infection. Frequency of donor-derived GC T_FH_ cells are shown in **F** (n = 6). **G-H**, Representative flow cytometry plots of IgD^lo^CD138^+^ plasma cells at 8 days post infection. Summary of the frequency of plasma cells are shown in **H** (n = 6). **I-J**, Summary of the frequency of ICOS^hi^CXCR5^+^ and Bcl-6^hi^CXCR5^+^ T_FH_ cells gated on adoptive cells identified by CD45.2 staining in each group at 8 days post infection (n = 6). **K**, LCMV-specific IgG concentration in serum from bone marrow chimeric mice at days 8 post-infection were measured by ELISA (n = 4 per group). P values are calculated using two-tailed Student’s t-test for paired comparisons or one-way ANOVA for multiple comparisons. *denotes p < 0.05, **denotes p < 0.01,***denotes p < 0.001 for the indicated comparison. n.s. = not significant.

We next evaluated the effect of TOX2 on GC B cell responses. TOX2 overexpression rescued the frequencies of GL-7⁺Fas⁺ GC B cells and IgD^lo^CD138^+^ plasma cells, and again, co-expression with Myc yielded a stronger rescue (Figures 4G–H, S7A–B). In bone marrow chimeras, TOX2 overexpression largely restored LCMV-specific IgG titers, whereas Myc alone did not. Notably, co-expression of TOX2 and Myc did further improve antigen-specific IgG production beyond the level achieved by TOX2 alone (Figure 4K). Collectively, these data demonstrate that NAT10 maintains Tox2 mRNA stability and that TOX2—but not Myc—is able to rescue the T_FH_ differentiation defect caused by NAT10 deficiency. This work reveals a previously unrecognized mechanism by which NAT10-mediated mRNA stability control regulates immune cell differentiation, providing new insight into the molecular pathways governing T_FH_ development and function.

### NAT10 and TOX2 promote maturation of the human T follicular helper cell response

To contextualize our findings, we integrated data from the Chinese Immune Multi-Omics Atlas (CIMA) provided by BGI Research, which includes a cohort of 428 adults who self-reported no active disease. The CIMA database consists of 6,484,974 high-quality single-cell RNA sequencing (scRNA-seq) data derived from peripheral blood mononuclear cells (PBMCs), with a median of 15,478 cells per individual. These PBMCs were categorized into five major cell types: innate lymphoid cells (ILCs) and natural killer (NK) cells, B cells, myeloid cells and hematopoietic stem and progenitor cells (HSPCs), CD4^+^ T cells, and CD8^+^ T cells. Upon analyzing CD4^+^ T cells via scRNA-seq, we observed that the proportion of CD4^+^ T follicular helper (T_FH_) cells was significantly elevated in females (Figure 5A). This observation suggests that females are more likely to generate robust antibody responses when exposed to external stimuli. This finding aligns with the well-established epidemiological observation that women are at higher risk for developing autoimmune diseases. Furthermore, we investigated the expression of NAT10 mRNA in CD4^+^ T cells and found that the proportion of T_FH_ cells was positively correlated with NAT10 expression. This indicates that NAT10 plays a crucial role in promoting the maturation of TFH cells in humans.

**Figure 5.**
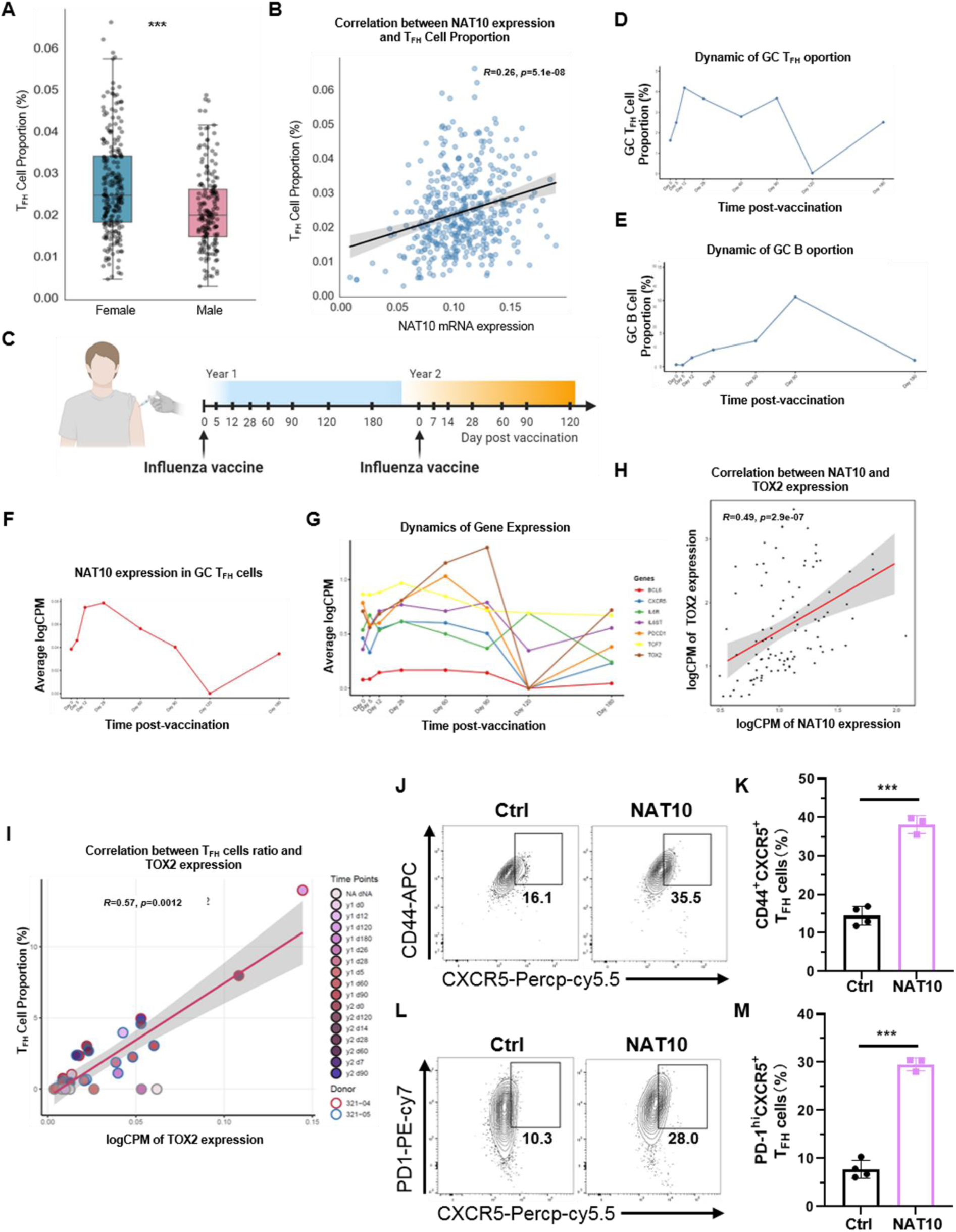
NAT10 and TOX2 promote human T follicular helper cell maturation. **A,** Box plot illustrating sex differences (female and male) in the proportions of T_FH_ cell in 428 Chinese individuals. **B,** Scatter plot comparing the NAT10 expression and T_FH_ cell proportion from 428 Chinese individuals. **C**, Schematic representation of the experimental approach used to profile blood and draining lymph node (LN) samples from human volunteers over a two-year period, following the administration of two influenza vaccines one year apart, to elucidate the evolution of the CD4^+^ T_FH_ cell response^[33]^. **D-E,** Frequency of GC T_FH_ and GC B cell in human volunteers over one year period. Shown as a percent of total CD4^+^ T cells or CD19^+^ B cells. **F**, The mRNA expression of NAT10 in GC T_FH_ cells populations from human volunteers over one year period. **G**, Scatter plot comparing the NAT10 expression and TOX2 expression in GC T_FH_ cell from human volunteers over two year period. Correlation analyses were performed using the Pearson correlation metric, and P values were computed using two-sided t test. **H**, The mRNA expression of TOX2 and key T_FH_-associated genes in GC T_FH_ cells populations from human volunteers over one year period. **I**, Scatter plot comparing the TOX2 expression and T_FH_ cell proportion from human volunteers over two year period. Correlation analyses were performed using the Pearson correlation metric, and P values were computed using two-sided t test. **J-M**, Flow cytometry analysis of CD44^+^CXCR5^+^ T_FH_ populations or PD-1^hi^CXCR5^+^ GC T_FH_ populations in human CD4^+^ T cells, which were activated *in vitro* under T_FH_ conditions and infected with Ctrl or NAT10 retrovirus, summary of the frequency of T_FH_ cells are shown in **K** and **M** (n = 3 per group).

To explore how NAT10 might regulate the maturation and differentiation of T_FH_ cells in response to vaccination, we extended our analysis by incorporating data from a longitudinal study of human volunteers who were followed over a two-year period following two influenza vaccinations^[33]^. By comparing the transcriptional profiles and cell populations of blood and draining lymph node (LN) samples, we were able to track the evolution of the CD4^+^ T_FH_ cell response over time (Figure 5C). Single-cell RNA sequencing (RNA-seq) analysis of peripheral blood mononuclear cells (PBMCs) and LN samples from the first year revealed that the frequency of germinal center (GC) T_FH_ cells among CD4^+^ T cells was initially low but expanded rapidly by Day 12 following vaccination, subsequently decreasing to baseline levels by Day 120 (Figure 5D). In parallel, the frequency of GC B cells among CD19^+^ cells increased gradually, reaching a peak at Day 90 post-vaccination (Figure 5E). Notably, the mRNA expression of NAT10 mirrored that of the GC T_FH_ cell population, showing a sharp increase at Day 12 post-vaccination, followed by a decline to baseline levels by Day 120 (Figure 5F). Additionally, the mRNA expression of key T_FH_-associated genes, including TOX2, exhibited a similar temporal pattern: an initial increase at Day 12 post-vaccination, followed by a decline to low levels by Day 120 (Figure 5G).We further investigated potential correlations between the mRNA expression of NAT10 and T_FH_-associated genes in GC T_FH_ cells, including TOX2, PDCD1, TCF7, IL6R, CXCR6 and BCL6. A significant positive correlation was observed between NAT10 and the expression of these T_FH_-associated genes at all time points, indicating a coordinated regulation of these genes during the immune response (Figure 5H and S8A-E). To determine whether NAT10 preferentially affects T_FH_ differentiation over other T helper subsets, we analyzed publicly available human PBMC scRNA-seq datasets. We found that the expression of Nat10 is positively correlated with the T_H_1 marker gene STAT1, but does not correlate with the expression of the T_H_2 marker gene GATA3 or the T_H_17 marker gene RORC (Figure S8F-G). This finding is consistent with the results observed following LCMV infection, where Nat10 knockout similarly affected different T cell subsets. Lastly, we examined the relationship between the expression of TOX2 and the proportion of T_FH_ cells. A significant positive correlation was found between the ratio of T_FH_ cells and TOX2 expression at all time points, further supporting the role of NAT10 and TOX2 in regulating T_FH_ cell dynamics (Figure 5I).

To directly demonstrate that NAT10 promotes T_FH_ cell differentiation, we isolated CD4^+^ T cells from human PBMCs and induced their differentiation into T_FH_-like cells *in vitro*. First, we investigated the effect of NAT10 overexpression on TOX2 expression and found that it significantly upregulated TOX2 (Figure S9A-B). We acknowledge that the differentiation efficiency of T_FH_-like cells from human CD4^+^ T cells is relatively low, with approximately 15% of cells being CD44^+^CXCR5^+^. However, upon overexpression of NAT10, the proportion of CD44^+^CXCR5^+^ cells increased significantly to around 35% (Figure 5J-M). Moreover, we assessed the proportion of PD1^+^CXCR5^+^ T_FH_-like cells and found that NAT10 overexpression significantly enhanced this population. We also evaluated the mean fluorescence intensity of CXCR5, CD44, and PD1 proteins and observed that NAT10 overexpression markedly increased the expression levels of these proteins (Figure S9C-E). Collectively, these results provide strong evidence that both NAT10 and TOX2 contribute to the maturation and regulation of the human T follicular helper cell response following vaccination.

### Identification and characterization of R428 as a small-molecule inhibitor of NAT10

Given the critical role of NAT10 in driving T_FH_ responses, we aimed to identify a pharmacological inhibitor as a potential therapeutic tool. We expressed and purified recombinant human NAT10 protein (Figure 6A) and established a robust in vitro enzymatic activity assay, confirming that NAT10 is an ATP-dependent enzyme (Figure 6B–C). To explore how small molecules regulate epigenetics and gene expression in eukaryotic cells, we employed virtual screening—a method widely used in drug discovery to identify candidate molecules that bind to a target protein structure from large compound libraries. We selected human NAT10 as the primary target due to its available crystal structure and known regulatory metabolites, which could be used for validation^[34]^. Using a structure-based virtual screening approach against a library of 2,503 FDA-approved compounds, we carefully evaluated docking scores and performed individual inspection, ultimately identifying 56 metabolites as potential NAT10 binders (Figure 6D–E). To experimentally validate the virtual screening results, we conducted *in vitro* enzyme activity assays using purified human NAT10. The enzyme was incubated with a 21-bp RNA substrate in the presence of MgCl₂, ATP, acetyl-CoA, and individual test compounds. NAT10 activity was quantified by measuring ATP consumption. This approach confirmed R428 (also known as BMS-777607) as a top candidate predicted to bind NAT10. *In vitro* screening showed that R428 was one of the most potent inhibitors of NAT10 acetyltransferase activity among the tested compounds (Figure 6F). Structural analysis suggested that R428 occupies the acetyl-CoA binding pocket of NAT10, directly competing with the cofactor (Figure 6G). Dose-response assays determined the half-maximal inhibitory concentration (IC₅₀) of R428 against NAT10 to be in the low micromolar range (Figure 6H). Notably, R428 exhibited superior inhibitory potency compared to two previously reported NAT10 inhibitors, remodelin and fludarabine^[35, 36]^(Figure 6I), establishing it as a novel and potent chemical probe for NAT10.

**Figure 6.**
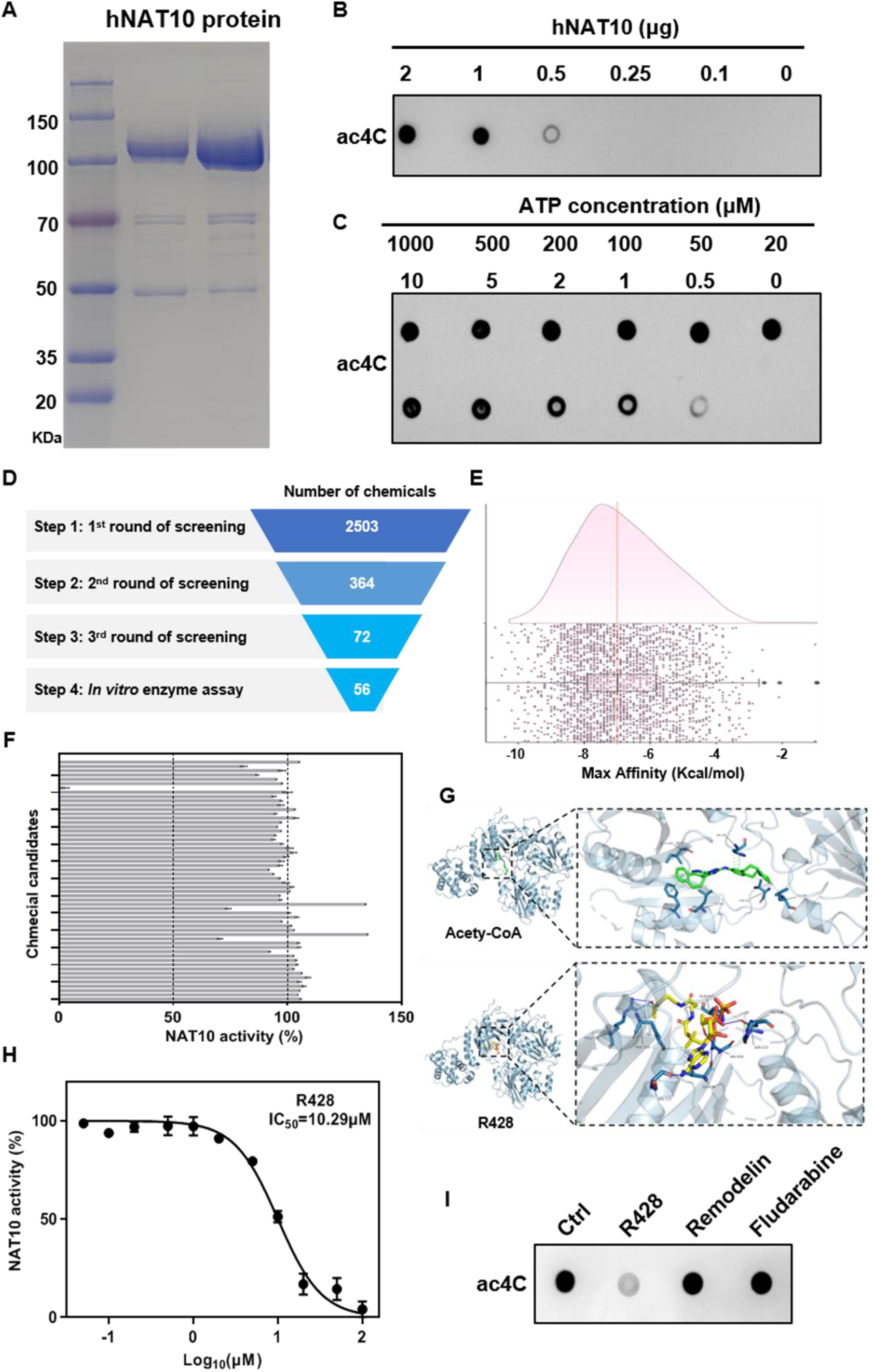
R428 is a potent NAT10 inhibor. **A,** Coomassie-stained gel of purified recombinant human NAT10 protein.. **B,** *In vitro* NAT10 enzymatic activity assay with increasing protein amounts, assessed by ac^4^C dot blot. **C**, *In vitro* activity assay with varying ATP concentrations. **D-E,** Schematic diagram of FDA-approved chemicals screening strategy via computational modeling of NAT10 binding. **F**, *In vitro* screen identifying R428 as a potent NAT10 inhibitor among 56 compounds.. **G**, The high-resolution 2.90 Å structure of NAT10 in complex with acety-CoA in PDB: 9J3C was used as the basis for modeling NAT10-R428 binding. **H**, Dose–response curve and half-maximal inhibitory concentration (IC_50_) of R428 were determined by the amount of ATP consumed. **I**, *In vitro* NAT10 activity assay showing inhibition by R428, but not by remodelin or fludarabine.

### Pharmacological inhibition of NAT10 alleviates pathology in allergic asthma

Finally, we evaluated the therapeutic efficacy of NAT10 inhibition in vivo using a T_FH_/GC-dependent disease model—ovalbumin (OVA)-induced allergic asthma^[21]^(Figure 6A). Oral administration of R428 (5 mg/kg/day) during the challenge phase significantly reduced the pathognomonic hallmarks of asthma. Treated mice exhibited lower levels of total IgE and IgG1 in the serum (Figure 7B-C) and reduced pro-inflammatory cytokines in lung homogenates (Figure 7D-F). Flow cytometric analysis of mediastinal lymph node (mLN) revealed that R428 treatment markedly attenuated the expansion of pathogenic CD44^+^CXCR5^+^ T_FH_ cells, PD-1^hi^CXCR5^+^ GC T_FH_ cells, and their associated IgD^lo^CD138^+^ plasma cells and GL-7^+^Fas^+^ GC B cells (Figure 7G-H and S10A-D). Histologically, R428 treatment resulted in substantial amelioration of lung pathology: reduced inflammatory cell infiltration (Figure 7I), decreased collagen deposition (Figure 7J), and attenuated goblet cell hyperplasia and mucus production (Figure 7K). These findings demonstrate that pharmacological targeting of NAT10 with R428 can effectively dampen dysregulated T_FH_ responses and mitigate immunopathology in a preclinical model of allergic disease.

**Figure 7.**
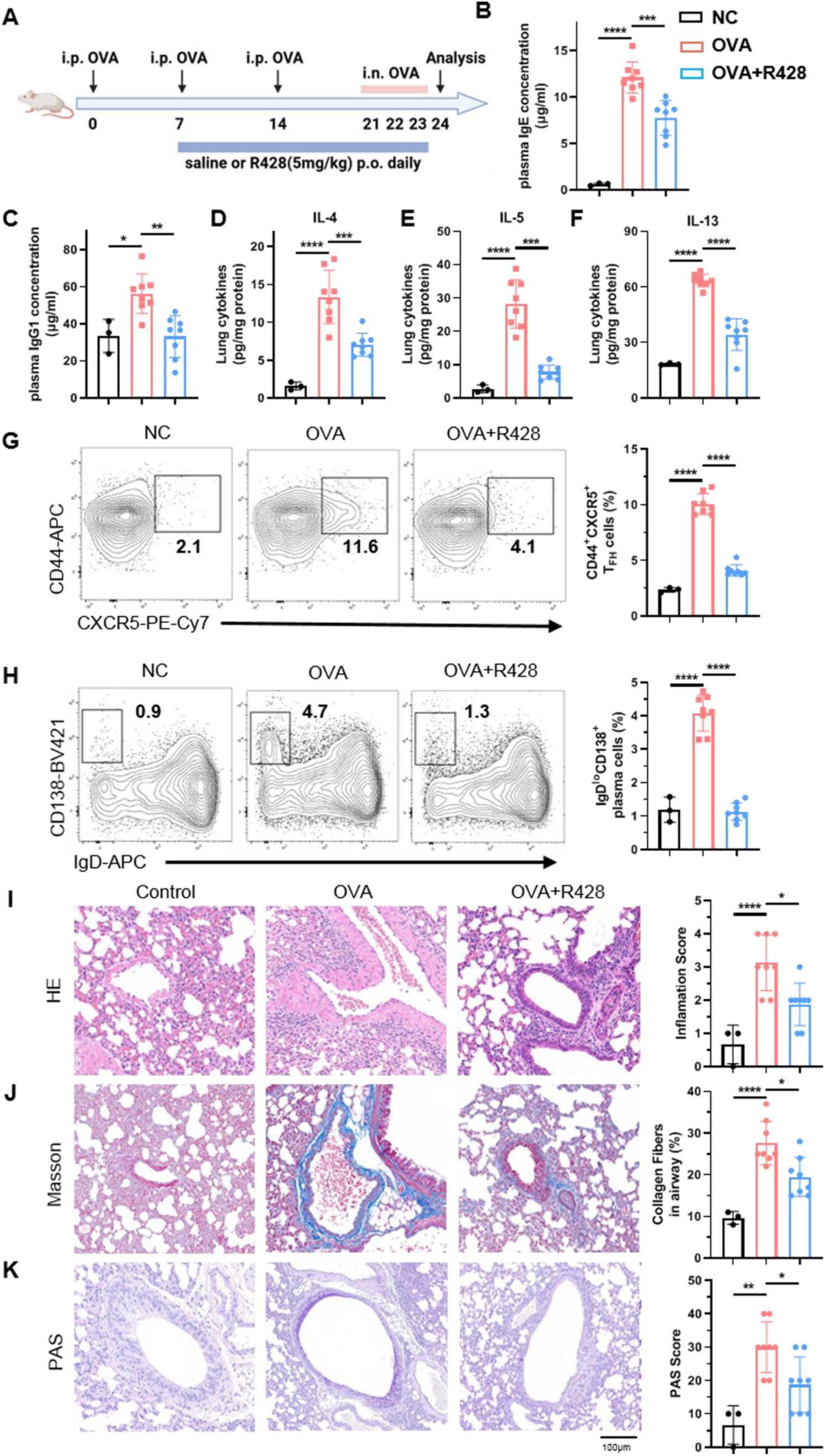
R428 treatment ameliorates allergic asthma by inhibiting T_FH_ responses. **A,** Schematic representation of ovalbumin (OVA)-induced allergic asthma model. Wild-type mice were primed with OVA and challenged with OVA intranasally. Saline or R428 (5mg/kg) orally administered to the mice once a day for 14 days. **B-C,** Total serum IgE and IgG1 levels at day 24. **D-F**, Cytokine levels in lung homogenates from mice with asthma treated with saline or R428 were analyzed by means of ELISA at day 24. **G**, Flow cytometry analysis and frequency of CD44^+^CXCR5^+^ T_FH_ cells, gated on mLN CD4^+^ T cells, from mice with asthma treated with saline or R428 at day 24 (n =3-8). **H**, Flow cytometry analysis and frequency of IgD^lo^CD138^+^ plasma cells in mLN from mice with asthma treated with saline or R428 at day 24 (n =3-8). **I**, The morphological changes of airway were observed by HE staining (left panel). The inflammatory infiltration was quantified by HE scores (right panel). **J**, Representative images of Masson’s staining of lung from mice with asthma treated with saline or R428 (left panel). The fibers was quantified by Masson’s staining (right panel). **K**, Representative images of PAS-stained lung sections from mice with asthma treated with saline or R428 (left panel). The goblet cell hyperplasia was determined by PAS scores (right panel). Scale bars: 100 µm. P values are calculated using two-tailed Student’s t-test for paired comparisons or one-way ANOVA for multiple comparisons. *denotes p < 0.05, **denotes p < 0.01,***denotes p < 0.001 and ****denotes p < 0.0001 for the indicated comparison.

## Discussion

T follicular helper (T_FH_) cells are essential for orchestrating germinal center (GC) responses and promoting long-lived humoral immunity. While transcriptional and epigenetic regulation of T_FH_ differentiation has been extensively studied, post-transcriptional mechanisms, particularly RNA modifications, remain largely unexplored in this lineage. Here, we identify Nat10 as a critical regulator of T_FH_ cell development via its ability to install ac⁴C RNA modifications. Our study demonstrates that T cell-specific Nat10 deletion leads to impaired T_FH_ cell differentiation, defective GC B cell responses during viral infection, and transcriptional downregulation of key T_FH_ -associated genes.

Through transcriptome and ac⁴C-RIP sequencing, we identified TOX2, a known transcription factor enriched in T_FH_ cells, as a direct ac⁴C-modified target of Nat10^[32, 37]^. Nat10 deficiency resulted in reduced ac⁴C modification and destabilization of TOX2 mRNA, leading to decreased TOX2 expression. Importantly, ectopic expression of TOX2 in Nat10-deficient T cells rescued T_FH_ differentiation and restored expression of signature molecules including Bcl6 and CXCR5. These findings define a previously unrecognized Nat10-ac⁴C-TOX2 regulatory axis essential for T_FH_ lineage specification. Nat10 could regulate ac^4^C modification on mRNAs, rRNAs, and tRNAs. Although we observed TOX2 as a direct ac⁴C-modified target of Nat10, we can not rule out the effect of rRNA/tRNA effects.

Our results add a new dimension to the regulatory network of T_FH_ differentiation by placing RNA acetylation alongside transcriptional and epigenetic programs. Prior studies have shown that TOX2 supports Tfh gene expression and represses alternate CD4^+^ T cell fates^[32]^, but upstream regulatory mechanisms of TOX2 remain poorly understood. Our findings now implicate ac⁴C as a functional post-transcriptional layer controlling TOX2 stability, aligning with recent reports of ac⁴C promoting mRNA stability and translation. Notably, Nat10 has been implicated in hematopoietic stem cell maintenance and cancer biology, but this study extends its function to adaptive immunity.

Furthermore, it opens the door to therapeutic intervention. Dysregulated T_FH_ cells and excessive GC reactions are central to a wide range of autoimmune diseases, allergies, and lymphomas. Our identification of R428 as a potent, specific inhibitor of NAT10’s acetyltransferase activity provides a novel pharmacological tool to test whether modulating this axis has therapeutic value. The efficacy of R428 in the OVA-induced asthma model is particularly compelling. By inhibiting NAT10, R428 treatment successfully dampened the pathogenic T_FH_

/GC/plasma cell axis, leading to reduced antibody titers and a dramatic alleviation of lung inflammation and remodeling. This proof-of-concept demonstrates that targeting epitranscriptomic pathways in vivo is feasible and can yield significant therapeutic benefits in a T_FH_-dependent disease. R428, with its known pharmacokinetic profile, could serve as a lead compound for developing next-generation NAT10 inhibitors.

While our study defines a clear NAT10-ac^4^C-TOX2 axis, several questions remain. First, is the regulation of Tox2 mRNA sufficient to explain all aspects of the NAT10-deficient phenotype? Our rescue experiments strongly suggest it is the major effector, but we cannot exclude contributions from other ac^4^C-modified transcripts. A comprehensive identification of the NAT10-regulated “ac^4^C epitranscriptome” in T_FH_ cells, coupled with rigorous genetic tests of individual targets, will map the full regulatory network. Second, what are the upstream signals that regulate NAT10 activity or expression in T cells during an immune response? Understanding whether NAT10 is instructed by T cell receptor strength, cytokine signals (e.g., IL-6, IL-21), or co-stimulation will place this epitranscriptomic regulator within the established signaling pathways of T_FH_ differentiation. Third, R428, previously reported as an inhibitor of AXL^[38]^, raises several questions regarding its role in Tfh differentiation and maturation. Whether AXL plays a role in the differentiation and maturation of Tfh cells, and whether the inhibitory effect of R428 on this process relies solely on its suppression of nat10 or is partially mediated through nat10, as well as the specific function of AXL in this pathway, all require further investigation. Future work should explore conditional or acute inhibition strategies, and investigate the role of NAT10 in other contexts where T_FH_ cells are important, such as anti-tumor immunity or chronic viral infections. Finally, the structural basis of R428 inhibition provides a roadmap for rational drug design to improve potency and selectivity.

In summary, we have discovered that the RNA acetyltransferase NAT10 functions as a pivotal epitranscriptomic regulator of T cell fate. By installing ac^4^C modifications on key mRNAs—most notably Tox2—NAT10 ensures their stable and efficient translation, thereby powering the gene expression program required for T_FH_ cell differentiation and effective humoral immunity. This pathway is conserved in humans and linked to vaccine responses. The identification of NAT10 as a druggable target and the efficacy of its inhibitor R428 in a disease model transform a fundamental biological insight into a promising therapeutic strategy. Our work not only expands the molecular toolkit controlling helper T cell differentiation but also establishes the epitranscriptome as a viable and potent layer for immune modulation, with broad implications for treating antibody-mediated diseases.

## Methods

### Mice

In this study, six- to eight-week-old C57BL/6J wild-type (WT) mice were obtained from SPF Biotechnology (Beijing, China). The conditional knockout mice harboring the FLOX allele were generously provided by Dr. Y. Lu (Shanghai Jiao Tong University, Shanghai, China). To facilitate the generation of the *Nat10*^fl/fl^*Cd4*^Cre^ mice, the *Nat10*^fl/fl^ mice were crossed with *Cd4*^Cre^ transgenic mice, which were acquired from Cyagen Biosciences (Guangzhou, China). All animals were housed and handled according to the institutional guidelines in pathogen-free facilities at Cyagen Biosciences and the Shanghai Public Health Clinical Center (Shanghai, China). The mice were maintained under specific pathogen-free (SPF) conditions, which were rigorously monitored to ensure the health and well-being of the animals throughout the course of the study. They were fed a standard diet of SPF-certified chow (SPF-F02-002). The animal protocols used in this study were approved by the Institutional Animal Care and Use Committee of the Shanghai Public Health Clinical Center (Approval No. 2023-A053-01), ensuring compliance with ethical standards for animal research.

### CD4^+^ T cell isolation and culture

Total CD4^+^ T cells were isolated using the MojoSort Mouse CD4 T Cell Isolation Kit (Biolegend), following the manufacturer’s instructions. To generate in vitro-activated CD4^+^ T cells, CD4^+^ T cells were stimulated with plate-bound anti-CD3ε (Biolegend, clone 2C11; 5 μg ml–1) and soluble anti-CD28 (eBioscience, clone 37.51; 0.5 μg ml–1) in the presence of IL-2 (PeproTech; 100 U ml–1). The culture medium consisted of RPMI-1640 supplemented with 10% fetal bovine serum (Gibco), 2 mM L-glutamine (Gibco), 1 mM sodium pyruvate (Sigma-Aldrich), 10 mM HEPES (Sigma-Aldrich), and 50 μM β-mercaptoethanol (Sigma-Aldrich). For mouse T_FH_ induction, naive CD4^+^ T cells were purified using the MojoSort Mouse CD4 Naïve T Cell Isolation Kit (Biolegend) according to the manufacturer’s instructions. The naive CD4^+^ T cells were stimulated with plate-bound anti-CD3ε (Biolegend, clone 2C11; 5 μg/ml) and soluble anti-CD28 (eBioscience, clone 37.51; 1 μg/ml) under neutral conditions anti-IFNγ (Bioxcell,10 μg/ml) and anti-IL4 (Tonbo, 10 μg/ml)) for 20 hours. Following this, the cells were infected with retrovirus in the presence of polybrene (8 μg/ml) using spin infection (2000 rpm, 32°C) for 90 minutes, after which fresh medium was changed 4 hours post-infection. One day after infection, GFP^+^ cells were sorted by FACS and cultured in medium containing anti-IFNγ antibody (Bioxcell,10 μg/ml), anti-IL-4 antibody (Bioxcell, 10 μg/ml), anti-TGF-β (Bioxcell, 20 μg/ml), IL-6 (PeproTech, 10 ng/ml), and IL-21 (Sino Biological, 10 ng/ml,). The culture medium used contained RPMI-1640, 10% fetal bovine serum (Gibco), 2 mM L-glutamine (Gibco), 1 mM sodium pyruvate (Sigma-Aldrich), 10 mM HEPES (Sigma-Aldrich), and 50 μM β-mercaptoethanol (Sigma-Aldrich). For human T_FH_ induction, human naïve CD4^+^ T cells were purified using the MojoSort Human CD4 Naïve T Cell Isolation Kit (Biolegend) according to the manufacturer’s instructions. The human naive CD4^+^ T cells were stimulated with plate-bound anti-human CD3ε (Biolegend, clone OKT3; 5 μg/ml) and soluble anti-human CD28 (Biolegend, clone CD28.2; 1 μg/ml) in RPMI 1640 (Sigma-Aldrich) supplemented with 10% fetal bovine serum (FBS) and penicillin and streptomycin (100 U/ml; Thermo Fisher Scientific) at 48-well plates (1 ml of medium per well). The culture medium was additionally supplemented with IL-6 (10 ng/ml), transforming growth factor–β (TGF-β; 5 ng/ml) and interleukin-23 (IL-23; 25 ng/ml; both from R&D Systems). To avoid medium exhaustion-dependent artificial effects on T_FH_ polarization, culture medium supplemented with IL-6, TGF-β and IL-23 was refreshed once on day 3 after stimulation.

### Plasmid construction, virus production and virus infection

The TOX2 or Myc overexpression plasmids were constructed using retroviral vectors MIGR1-IRES-GFP (a gift from H. Hu, Sichuan University) via homologous recombination. Plasmid was verified by Sanger sequencing (Genewiz) and propagated in DH5α competent cells (Tsingke). To produce retroviral particles, 293T cells were transfected with the plasmids using Lipo6000 (Beyotime). Forty-eight hours post-transfection, supernatants containing 10 µg/mL Polybrene (Beyotime) were collected, filtered, and mixed with HSCs. The mixtures were then centrifuged at 800 × g for 2 hours at 33 °C, followed by incubation for 3 hours at 37 °C. The cells were subsequently transferred to fresh culture medium or collected for further processing. Infected cells were identified based on fluorescence and analyzed by flow cytometry.

### LCMV infection

The LCMV Armstrong strain was generously provided by Dr. L. Ye (Army Medical University, Chongqing, China). Virus propagation and titration were performed as previously described^[39]^. Briefly, the LCMV-Armstrong strain was propagated in BHK-21 cells, and viral titers were determined by plaque assays to ensure accurate quantification. To establish an acute LCMV infection, mice were intraperitoneally injected with 2 × 10^5 plaque-forming units (p.f.u.) of LCMV-Armstrong, as described in prior studies. For the generation of bone marrow chimeric mice,

LCMV infection was conducted following an 8-week reconstitution period, allowing sufficient time for the donor-derived hematopoietic cells to fully repopulate the recipient’s immune system.

### Flow cytometry and antibodies

Single-cell suspensions of spleens were used for flow cytometry analysis or cell sorting. Surface staining was performed in PBS containing 1% FBS. The antibodies and reagents used for flow cytometry staining are listed as: anti-CD44 (IM7; BioLegend), anti-CXCR5 (L138D7; BioLegend), anti-CXCR5 (560617; BD Biosciences), anti-PD1 (29F.1A12; BioLegend), anti-GL-7 (561529; BD Biosciences), anti-Fas (561979; BD Biosciences), anti-ICOS (15F9; BioLegend), anti-Bcl-6 (7D1; BioLegend), anti-IgD (11-26c; Thermo Fisher Scientific), anti-CD138 (562610; BD Biosciences), anti-SLAM (TC15-12F12.2; BioLegend), anti-CD4 (RM4-5; BioLegend), anti-CD19 (1D3/CD19; BioLegend), anti-CD45.1 (A20; BioLegend), anti-CD45.2 (104; BioLegend), anti-CD4 (GK1.5; BioLegend), anti-TOX2 (21162AP, ProteinTech), anti-NAT10 (ab194297, Abcam). Samples were acquired on a BD FACSLyric Cytometer. All data analysis was performed using FlowJo 10.4 software (FlowJo). Sorting assays were performed using a FACS Aria Cytometer (BD Bioscience). Gating strategies are presented in Figure S5.

### Bone marrow chimeric mice

Bone marrow chimeric mouse models were generated as previously described ^[18]^. Briefly, *Nat10*^fl/fl^ and *Nat10*^fl/fl^*Cd4*^Cre^ donor mice (CD45.2^+^) were administered 5-fluorouracil (Sigma-Aldrich) intraperitoneally at a dose of 150 mg per kg body weight, prepared in PBS. After 24 hours, hematopoietic stem cells (HSCs) were enriched from bone marrow using a CD117^+^ selection kit (Stem Cell Technologies) and subsequently transduced with retroviral supernatants containing MIGR1 plasmids via spin infection, as previously outlined. During HSC transfection, recipient mice (CD45.1^+^) were sublethally irradiated (9 Gy). HSCs were resuspended at 5 × 10^6 cells per ml. Each recipient mouse was then intravenously injected with 0.2 ml of the cell suspension (1 million total cells). After 8 weeks reconstitution, recipient mice were infected with LCMV-Armstrong strain.

### RNA ac^4^C dot blot

Purified RNA diluted to the indicated concentrations and incubated at 95 °C for 3 minutes. Equal volumes of RNA samples were then transferred onto two separate nitrocellulose membranes. After air-drying for 20 minutes, the RNA was cross-linked to the membranes by UV irradiation (5 J, three times). One membrane was stained with methylene blue (0.02% in 0.3 M sodium acetate) and washed with double-distilled water for 1 hour to serve as a loading control. The second membrane was blocked with 5% bovine serum albumin and incubated overnight at 4 °C with anti-ac^4^C antibody (Abcam), followed by incubation with a secondary antibody (Proteintech) for 90 minutes at room temperature. Blot signals were detected using a ChemiScope 6100 (Clinx Science Instruments), and densitometric analysis was performed using ImageJ v.1.53a software.

### RIP–qPCR

The ac^4^C RIP assay was conducted using the GenSeq ac^4^C RIP kit (GS-ET-005, Cloudseq Biotech) according to the manufacturer’s instructions. Briefly, total RNA (100 µg) was randomly fragmented into 100- to 200-bp pieces and incubated with 5 µg of either anti-ac^4^C or IgG antibodies, along with magnetic beads. RNA bound to anti-ac^4^C was then purified, and enrichment of *Tox2* mRNA was assessed by qPCR following reverse transcription. For the NAT10 RIP assay, anti-ac^4^C was replaced with anti-NAT10, while all other steps remained unchanged. The primers used are provided in Supplementary Table 4.

### RNA degradation assay

T follicular helper (T_FH_) cells were isolated from *Nat10*^fl/fl^*Cd4*^Cre^ mice and their wild-type controls and cultured as described previously. And subsequently treated with 2.5 µg/ml actinomycin D for 0, 1, or 2 hours. Total cellular RNA was then extracted, and the relative transcriptional expression of Tox2 was quantified by qPCR following reverse transcription.

### ELISA

LCMV-specific antibody levels in serum were analyzed as previously described^[40]^. In brief, lysates of LCMV-infected BHK-21 cells were used as the substrate, and LCMV-specific IgG antibodies were titrated through serial dilutions of serum. Detection was performed using HRP-conjugated goat anti-mouse IgG antibodies (Cat. no. A90-131P-39; 1:5,000; Bethyl Laboratories).

### Polyribosome real-time PCR

T follicular helper (T_FH_) cells were isolated from *Nat10*^fl/fl^*Cd4*^Cre^ mice and their wild-type controls as previously described, and subsequently incubated with cycloheximide (100 µg/mL; MedChem Express) for 5 minutes. Cells were collected and washed with prechilled PBS containing 100 µg/mL cycloheximide. Following lysis, one-third of the lysate was retained as input, and the remaining lysate was gently layered onto a sucrose gradient buffer. Centrifugation was performed at 274,000 × g for 1.5 hours at 4 °C. Gradient fractions were collected using a Biocomp Piston Gradient Fractionator. RNase inhibitor (Beyotime) was added to the entire system at a final concentration of 1,000 U/mL. RNA was extracted from polyribosome fractions and input using the RNA Clean & Concentrator kit (Zymo Research) and analyzed by qPCR.

### acRIP–seq

The acRIP-Seq service was provided by CloudSeq Biotech. Total RNA was extracted from CD4^+^ cells sorted from LCMV infected *Nat10*^fl/fl^*Cd4*^Cre^ or Ctrl mice, with rRNA depletion achieved using a Ribo-zero kit (Illumina). ac^4^C immunoprecipitation (ac^4^C-IP) was performed according to the manufacturer’s instructions using a GenSeq ac^4^C-IP kit (GenSeq). In brief, RNA was fragmented into fragments smaller than 200 nucleotides using RNA fragmentation reagents. Anti-ac^4^C was conjugated to Protein A/G beads by incubation with gentle rotation at room temperature for 1 hour. Fragmented RNA was then incubated with the antibody-bound beads, rotating at 4 °C for 4 hours. Immunoprecipitated samples were treated with Proteinase K, and RNA was recovered via phenol/chloroform extraction. RNA libraries were subsequently constructed using the NEBNext Ultra II Directional RNA Library Prep kit (New England Biolabs) following the manufacturer’s guidelines. The quality of the libraries was assessed using an Agilent BioAnalyzer 2100 system, and sequencing was performed on the Illumina NovaSeq platform. Raw sequencing reads from the Illumina NovaSeq 6000 were quality-controlled by Q30 and processed with Cutadapt software (v1.9.3) to trim 3’ adaptors and remove low-quality alignments. The filtered reads were aligned to the reference genome using HISAT2 (v2.0.4). ac^4^C-enriched regions (peaks) were identified with MACS software, and differentially acetylated sites were determined with diffReps. Peaks that overlapped with exons were selected using in-house scripts. Genomic distributions of ac^4^C peaks were visualized with the Integrative Genomics Viewer (IGV). Motif analysis was conducted using the Discriminative Regular Expression Motif Elicitation software according to the default workflow. Meta-gene analysis of ac^4^C sites on mRNAs from T_FH_ cells was performed using the metaPlotR package.

### RNA-seq and data analysis

In this study, RNA sequencing services were provided by CloudSeq Biotech. To isolate T_H_1 and T_FH_ cells, splenocytes from *Nat10*^fl/fl^*Cd4*^Cre^ mice and control littermates at day 8 post-viral infection were surface stained, and CD44^+^SLAM^hi^ T_H_1 cells and CD44^+^SLAM^lo^ T_FH_ cells were sorted using flow cytometry, and total RNA was extracted. rRNA was removed using the NEBNext rRNA Depletion Kit (New England Biolabs) according to the manufacturer’s protocol. RNA libraries were constructed with the NEBNext Ultra II Directional RNA Library Prep Kit (New England Biolabs) following the provided instructions. After quality control and quantification on a BioAnalyzer 2100 system (Agilent Technologies), the libraries were sequenced on an Illumina NovaSeq 6000 platform with paired-end 150 bp reads. Following quality control using a Q30 threshold, raw data were further processed by trimming 3’ adaptors with cutadapt software (v1.9.3) to obtain high-quality clean reads. These reads were aligned to the reference genome (UCSC mm10) using HISAT2 software (v2.0.4). Raw counts were generated with HTSeq (v0.9.1), and differential expression analysis was performed using the edgeR package, with a |log2 (fold change)| ≥ 1 and a false discovery rate (FDR) ≤ 0.05. Gene ontology (GO) and pathway enrichment analyses were conducted using the clusterProfiler package. Gene set enrichment analysis (GSEA) was performed with the R package AUCell.

### RNA extraction and real-time qPCR

Total RNA was isolated using the RNA-Quick Purification Kit (ES Science). One microgram of total RNA was reverse transcribed with SYBR Premix Ex Taq II (Takara). The resulting cDNA samples were diluted and subsequently analyzed by quantitative real-time PCR using SYBR Premix Ex Taq II (Takara), according to the manufacturer’s guidelines, on a StepOnePlus Real-Time PCR System. Actin served as the endogenous normalization control. Primers were synthesized by Sangon Biotech, with the exact sequences provided in Supplementary Table 4.

### Western blotting

Cells were lysed on ice for 30 minutes in RIPA buffer (Beyotime), supplemented with protease and phosphatase inhibitors (Beyotime). The lysates were then centrifuged at 12,000g for 5 minutes at 4°C to collect the supernatant. Protein concentrations were determined using a bicinchoninic acid (BCA) assay (Beyotime) and adjusted to a final concentration of 1–2 μg/μl. Samples were mixed with 5× SDS–PAGE loading buffer (Beyotime), heated at 100°C for 10 minutes, separated by SDS–PAGE, and transferred to a PVDF membrane (Millipore). The membranes were blocked in 5% skim milk at room temperature for 1 hour, followed by incubation with the appropriate primary and secondary antibodies. Protein signals were visualized using a ChemiScope 6100 (Clinx Science Instruments).

### Analysis of data from the CIMA and Influenza vaccination datasets

The CIMA dataset was generously provided by the Liu group at BGI Research, and the influenza vaccination datasets were obtained from a previously published study^[33]^. To analyze the scRNA-seq data, we employed a comprehensive workflow built upon established computational tools. These tools include Scrublet (version 0.2.3)^[41]^, which was used for cell doublet detection; Scanpy (version 1.9.3)^[42]^, a scalable Python library for the analysis of large single-cell RNA-seq datasets; and COSG (version 1.0.1)^[43]^, which facilitates the identification of cell-specific gene signatures. This robust computational pipeline was systematically applied to all scRNA-seq samples, encompassing both the CIMA cohort and the influenza vaccination datasets. The integration of these tools enabled us to perform rigorous quality control, cell type annotation, and downstream analyses to ensure the reliability and reproducibility of our findings across both datasets.

### Asthma Model Induction and Drug Treatment

Mice were sensitized by intraperitoneal injection of 50 μg ovalbumin (OVA) adsorbed to 2 mg aluminium hydroxide (Alum) in a 200 μL volume on days 0, 7, and 14. From day 7, the treatment group received the NAT10 inhibitor R428 (5 mg/kg in 50% PEG300/saline vehicle) via daily oral gavage until the endpoint of the experiment; control mice received vehicle only. On days 21-23, mice were challenged intranasally with 30 μL of 3% OVA solution under isoflurane anesthesia to induce allergic airway inflammation.

### In Vitro NAT10 Enzymatic Activity Assay

Recombinant human NAT10 protein was expressed in Escherichia coli BL21(DE3) and purified via affinity chromatography followed by ion exchange. A single-stranded RNA oligonucleotide (5’-ACAAGGUUUCCGUAGGUGAAC-3’) was synthesized as the substrate. The enzymatic reaction was performed in a 20 μL system containing 50 mM Tris-HCl (pH 7.5), 150 mM KCl, 5 mM MgCl₂, 1 mM DTT, 100 μM ATP, 500 μM acetyl-CoA, 1 μM RNA substrate, and purified NAT10. Test compounds were included as specified. Reactions were incubated at 37 °C for 3 hours with constant shaking at 700 rpm. NAT10 activity was quantified by measuring the production of ac⁴C via dot-blot analysis using an anti-ac⁴C antibody, or by monitoring ATP consumption.

### Histological Analysis

Lung tissues from mice were fixed in 10% neutral buffered formalin, embedded in paraffin, and sectioned at 4 μm. Following deparaffinization and rehydration, sections were subjected to histological examination. Inflammation was assessed by hematoxylin and eosin (HE) staining and scored based on established criteria^[44, 45]^. Collagen deposition was evaluated using Masson’s trichrome staining, with the mean percentage area quantified as previously described^[46]^. Goblet cell hyperplasia was examined via periodic acid–Schiff (PAS) staining according to the method of Padrid et al^[47]^.

### Statistical analysis

All experiments were conducted with a minimum of three biological replicates. Statistical analyses were performed with two-tailed Student’s t-test for paired comparisons or two-way ANOVA for multiple comparisons. Data shown (if not pointed out) represents the results obtained from triplicate independent experiments with mean +/- standard deviation (S.D.) or standard error of the mean (SEM). The values of p < 0.05 were considered statistically significant.

## Author contributions

D. Wu and Z-L. Cheng. conceived and supervised the project; Z-L. Cheng. designed and performed most of the experiments; Z-L. Cheng. and Z. Lin performed RNA-seq; Z. Lin. helped to breed *Nat10*^fl/fl^*Cd4*^Cre^ mice; Z-L. Cheng. executed ac^4^C RIP-seq and RNA-seq experiments; Z. Lin analyzed the RNA-seq and ac^4^C RIP-seq data. Z-L. Cheng. wrote the manuscript with the revision by all authors. D. Wu conceived experiments and supervised the study.

## Acknowledgements

We are grateful to the following researchers for providing valuable resources: Shuyang Yu (China Agricultural University) and Hongbo Hu (Sichuan University) for the MIGR1 plasmid; Yan Lu (Shanghai Jiao Tong University) for the *Nat10*^fl/fl^ mice; Caiwen Duan (Shanghai Jiao Tong University) for technical assistance; and Lilin Ye (Third Military Medical University) for the LCMV Armstrong virus. This work was supported by grants from the National Natural Science Foundation of China (82273188, 82570017) to D.W. and (32570687) to Z-L. Cheng; the Special Fund of the Science and Technology Commission of Shanghai Municipality (20244Z0013); the Shanghai Dongfang Top Talents Program (to D.W.); the Shanghai Dongfang Youth Talents Program (to Z-L. Cheng); and the Clinical Research Program of Zhongshan Hospital, Fudan University (2020ZSLC07) (to D.W.).

## Extended Data Figure Legends

**Figure S1.**
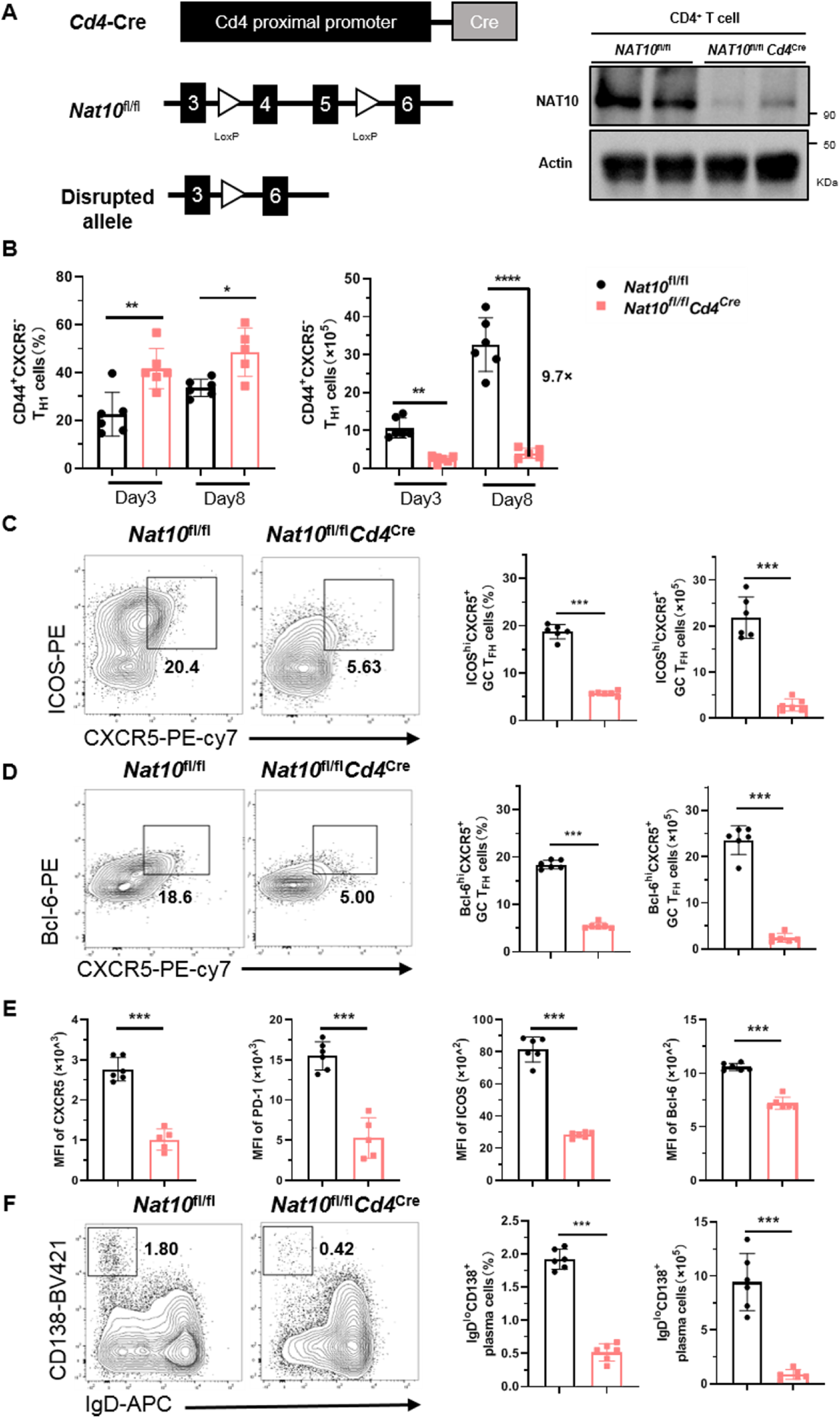
NAT10 is essential for T_FH_ differentiation and humoral immunity. **A**, *Cd4*-Cre mice were crossed with *Nat10*^fl/fl^ strain to generate conditional knockout mice. Recombination resulted in the excision of exons 4 and 5 of the *Nat10* gene, flanked by LoxP sites, leading to inactivation of NAT10 (left). Verification of *Nat10* depletion in CD4^+^ T cells from *Nat10*^fl/fl^ and *Nat10*^fl/fl^*Cd4*^Cre^ mice, as determined by western-blot (right). **B**, A summary of the frequency and cell numbers of CD44^+^CXCR5^-^ T_H_1 cells at days 3 and day 8 post-infection (n = 6). **C**, Flow cytometry analysis of ICOS^hi^CXCR5^+^ T_FH_ cells, gated on splenic CD4^+^ T cells, from *Nat10*^fl/fl^and *Nat10*^fl/fl^*Cd4*^Cre^ mice at day 8 post-infection (left). A summary of the frequency and cell numbers of indicated cell subsets at day 8 post-infection (n = 6) (right). **D**, Flow cytometry analysis of Bcl-6^hi^CXCR5^+^ T_FH_ cells, gated on splenic CD4^+^ T cells, from *Nat10*^fl/fl^and *Nat10*^fl/fl^*Cd4*^Cre^ mice at day 8 post-infection (left). A summary of the frequency and cell numbers of indicated cell subsets at day 8 post-infection (n = 6) (right). **E**, Quantification of the MFIs of CXCR5, PD-1, ICOS, and Bcl-6 on CD44^+^CXCR5^+^ T_FH_ cells at day 8 post-infection (n = 6). **F**, Flow cytometry analysis of IgD^lo^CD138^+^ plasma cells from *Nat10*^fl/fl^ and *Nat10*^fl/fl^*Cd4*^Cre^ mice at day 8 post-infection (left). A summary of the frequency and cell numbers of indicated cell subsets at days 8 post-infection (n = 6) (right). P values are calculated using two-tailed Student’s t-test for paired comparisons or one-way ANOVA for multiple comparisons. ***denotes p < 0.001.

**Figure S2.**
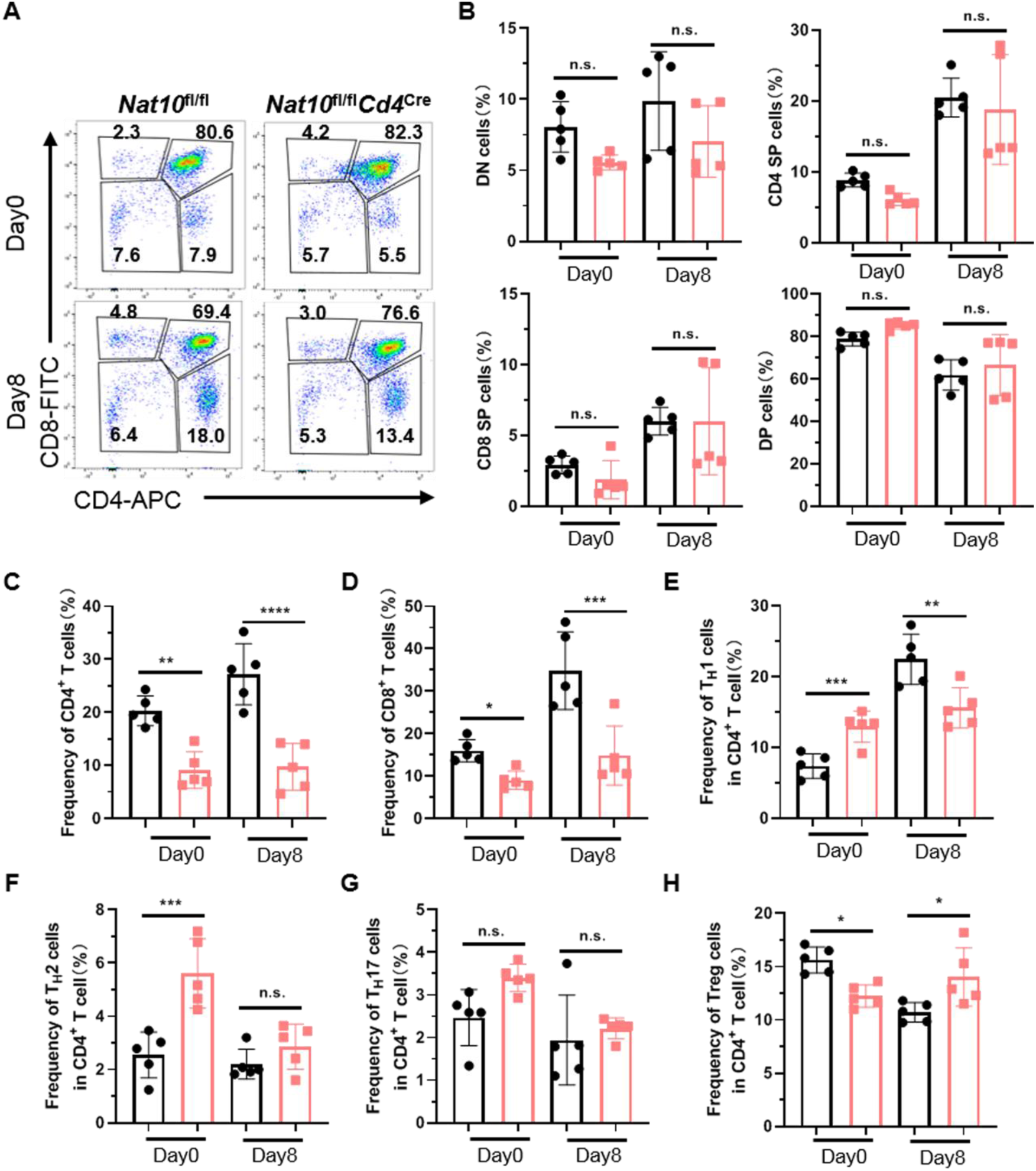
NAT10 knockout results in alterations in the proportions of various T cell subsets. **A-B**, Flow cytometry analysis of Thymic double-negative (DN), double-positive (DP) and CD4^+^ and CD8^+^ single-positive (SP) T cells, from *Nat10*^fl/fl^ and *Nat10*^fl/fl^*Cd4*^Cre^ mice at day 8 post-infection. A summary of the frequency of indicated cell subsets at day 8 post-infection (n = 5). **C-H**, The frequency of indicated cell subsets in spleen at day 8 post-infection (n = 5). P values are calculated using two-tailed Student’s t-test for paired comparisons or one-way ANOVA for multiple comparisons. *denotes p < 0.05, **denotes p < 0.01,***denotes p < 0.001 and ****denotes p < 0.0001 for the indicated comparison. n.s. = not significant.

**Figure S3.**
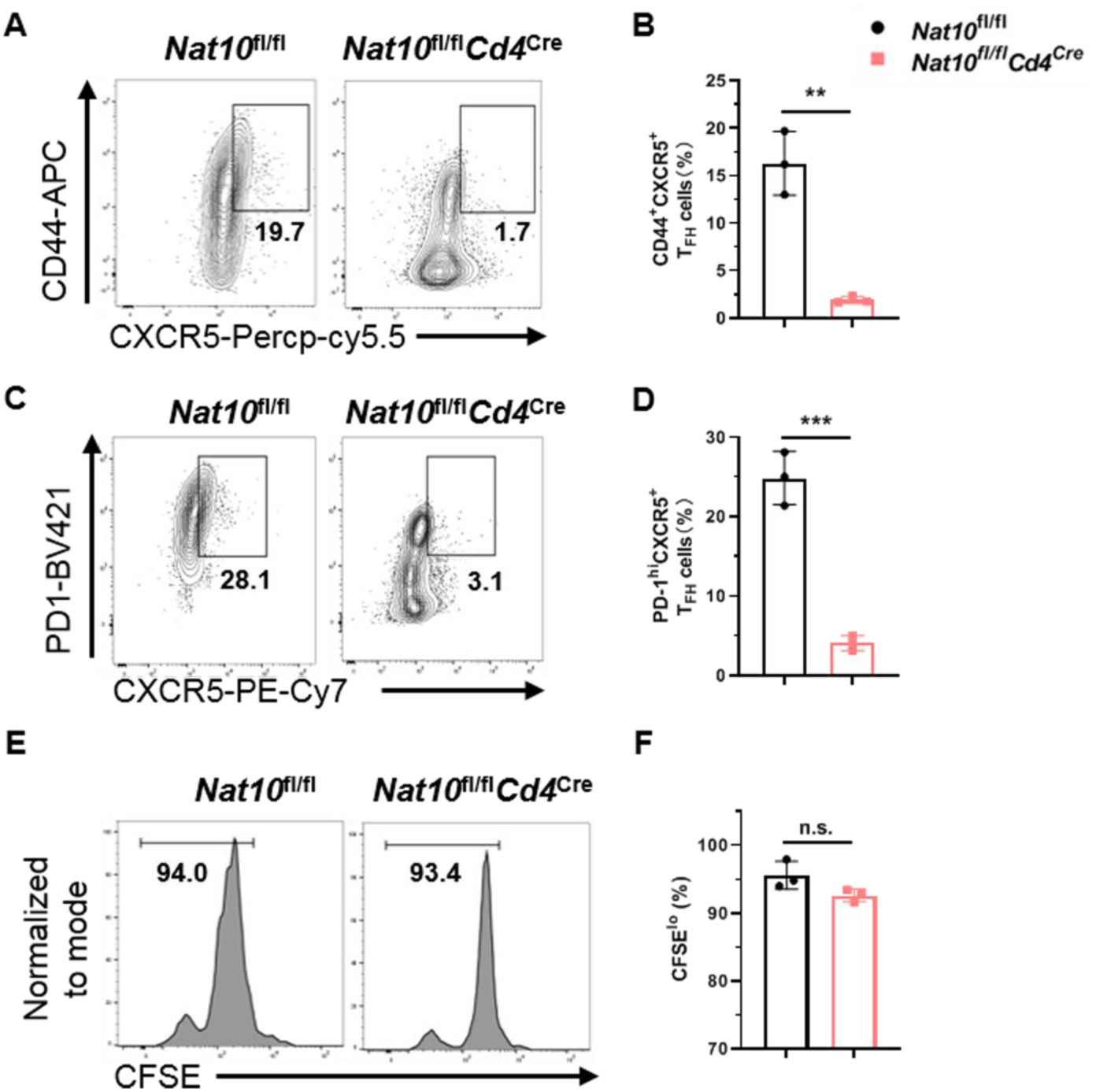
NAT10 deletion impairs T_FH_ differentiation *in vitro*. **A-D,** Flow cytometry analysis of CD44^+^CXCR5^+^ T_FH_ populations and PD-1^hi^CXCR5^+^ GC T_FH_ populations in mouse CD4^+^ T cells from *Nat10*^fl/fl^ and *Nat10*^fl/fl^*Cd4*^Cre^ mice, which were activated *in vitro* under T_FH_ conditions. The summary of the frequency of T_FH_ cells is shown in **B** and **D** (n = 3). **E-F**, Representative plots of CFSE staining in T_FH_ populations which were activated *in vitro* under T_FH_ conditions for 72 h. Bar graphs showing the percentages of CFSE^lo^ T_FH_ cells after 72 h of stimulation. P values are calculated using two-tailed Student’s t-test for paired comparisons. **denotes p < 0.01 and ***denotes p < 0.001 for the indicated comparison. n.s. = not significant.

**Figure S4.**
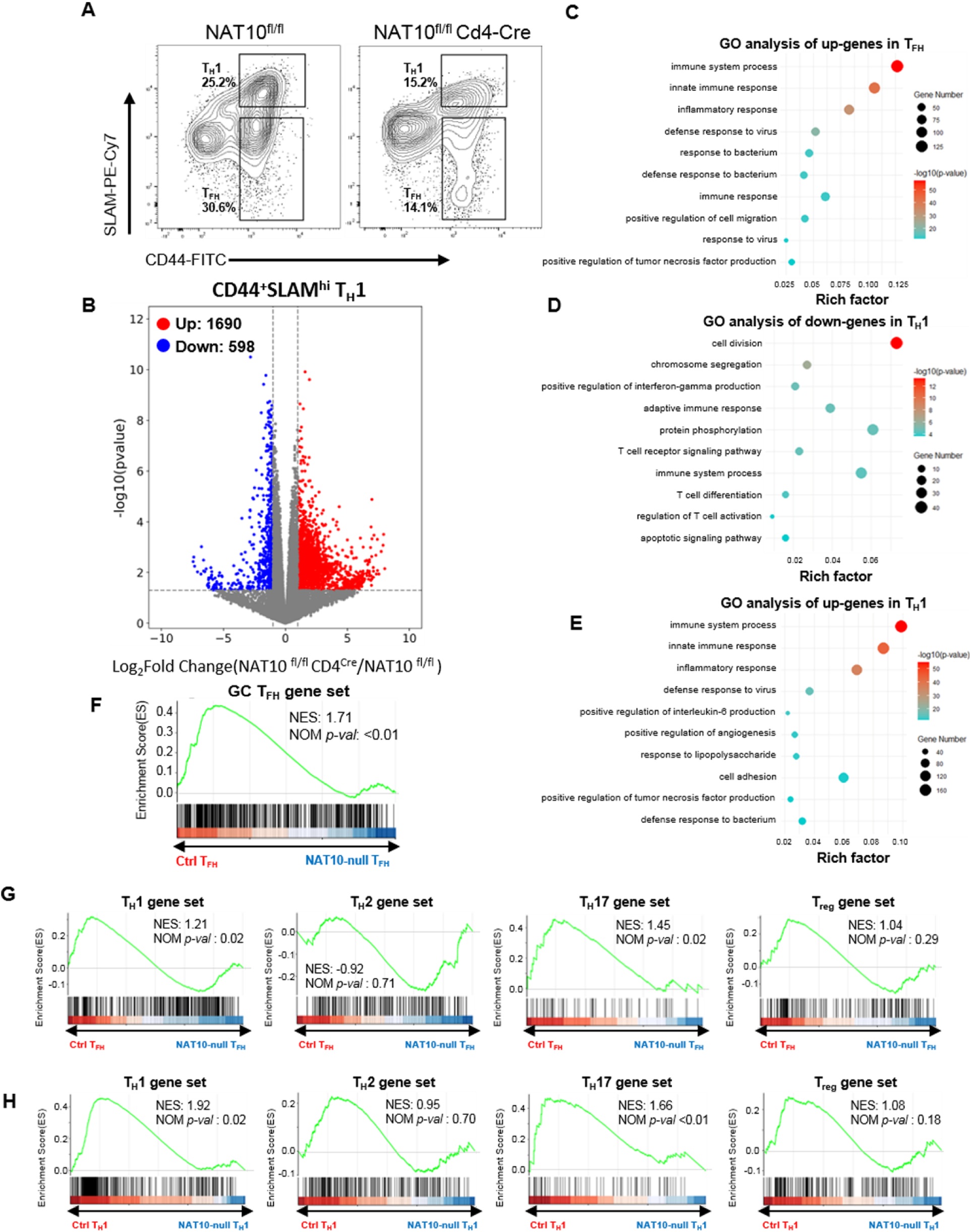
Transcriptional alterations in NAT10-deficient T_H_1 and T_FH_ cells. **A**, Flow cytometry analysis of CD44^+^SLAM^hi^ T_H_1 cells and CD44^+^SLAM^lo^ T_FH_ cells from *Nat10*^fl/fl^ and *Nat10*^fl/fl^*Cd4*^Cre^ mice at day 8 post-viral infection. **B**, Volcano plot showing differentially expressed genes between *Nat10*^fl/fl^ and *Nat10*^fl/fl^*Cd4*^Cre^ T_H_1 cells at day 8 post-viral infection. All differentially expressed genes with P<0.05 and fold change>2 are highlighted in red (up-regulation) and blue (down-regulation). **C**, Top GO enrichment analysis of the up-regulated genes in *Nat10*^fl/fl^*Cd4*^Cre^ T_FH_ cells compared with *Nat10*^fl/fl^ T_FH_ cells. **D**, Top GO enrichment analysis of the down-regulated genes in *Nat10*^fl/fl^*Cd4*^Cre^ T_H_1 cells compared with *Nat10*^fl/fl^ T_H_1 cells. **E**, Top GO enrichment analysis of the up-regulated genes in *Nat10*^fl/fl^*Cd4*^Cre^ T_H_1 cells compared with *Nat10*^fl/fl^ T_H_1 cells. **F**, Gene set enrichment analysis (GSEA) of GC TFH gene set in *Nat10*^fl/fl^*Cd4*^Cre^ T_FH_ cells relative to expression in *Nat10*^fl/fl^ T_FH_ cells. **G,** GSEA analysis of TH1, TH2, TH17, and Treg-associated gene sets in *Nat10*^fl/fl^*Cd4*^Cre^ T_FH_ cells relative to expression in *Nat10*^fl/fl^ T_FH_ cells. **H,** GSEA analysis of TH1, TH2, TH17, and Treg-associated gene sets in *Nat10*^fl/fl^*Cd4*^Cre^ T_H_1 cells relative to expression in *Nat10*^fl/fl^ T_H_1 cells.

**Figure S5.**
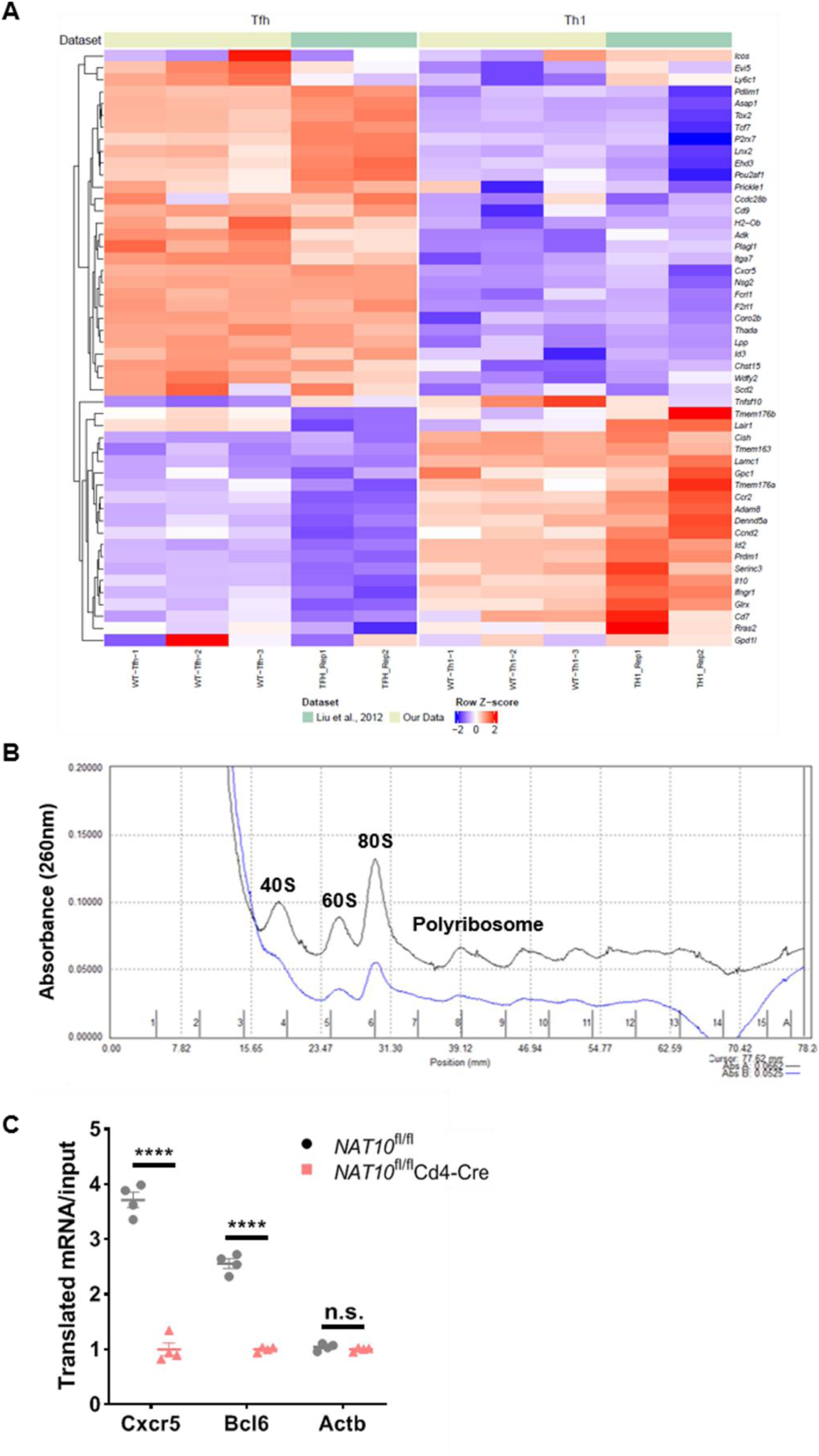
Altered gene expression and ribosome occupancy in NAT10-deficient T cells. **A**, Heatmaps representation of T_FH_ and T_H_1 associated genes in the CD44^+^SLAM^lo^ and CD44^+^SLAM^hi^ T cells compared with T_FH_ cells (CD4^+^CD44^+^CXCR5^+^BCL6^+^) and T_H_1 cells (CD4^+^CD44^+^CXCR5^-^BCL6^-^) from previous study. **B**, Representative ribosome profiles were obtained from ribosomal extracts prepared from T_FH_ cells in the presence of cycloheximide. The extracts were fractionated using a 5–50% sucrose gradient and analyzed with a pump-syringe apparatus coupled to an ultraviolet detector.. **C,** Ribosome occupancy of *Cxcr5*, *Bcl6* and control *Actb* mRNAs in *Nat10*^fl/fl^ and *Nat10*^fl/fl^*Cd4*^Cre^ T_FH_ cells. (n =4). P values are calculated using two-tailed Student’s t-test for paired comparisons or one-way ANOVA for multiple comparisons. ****denotes p < 0.0001 for the indicated comparison. n.s. = not significant.

**Figure S6.**
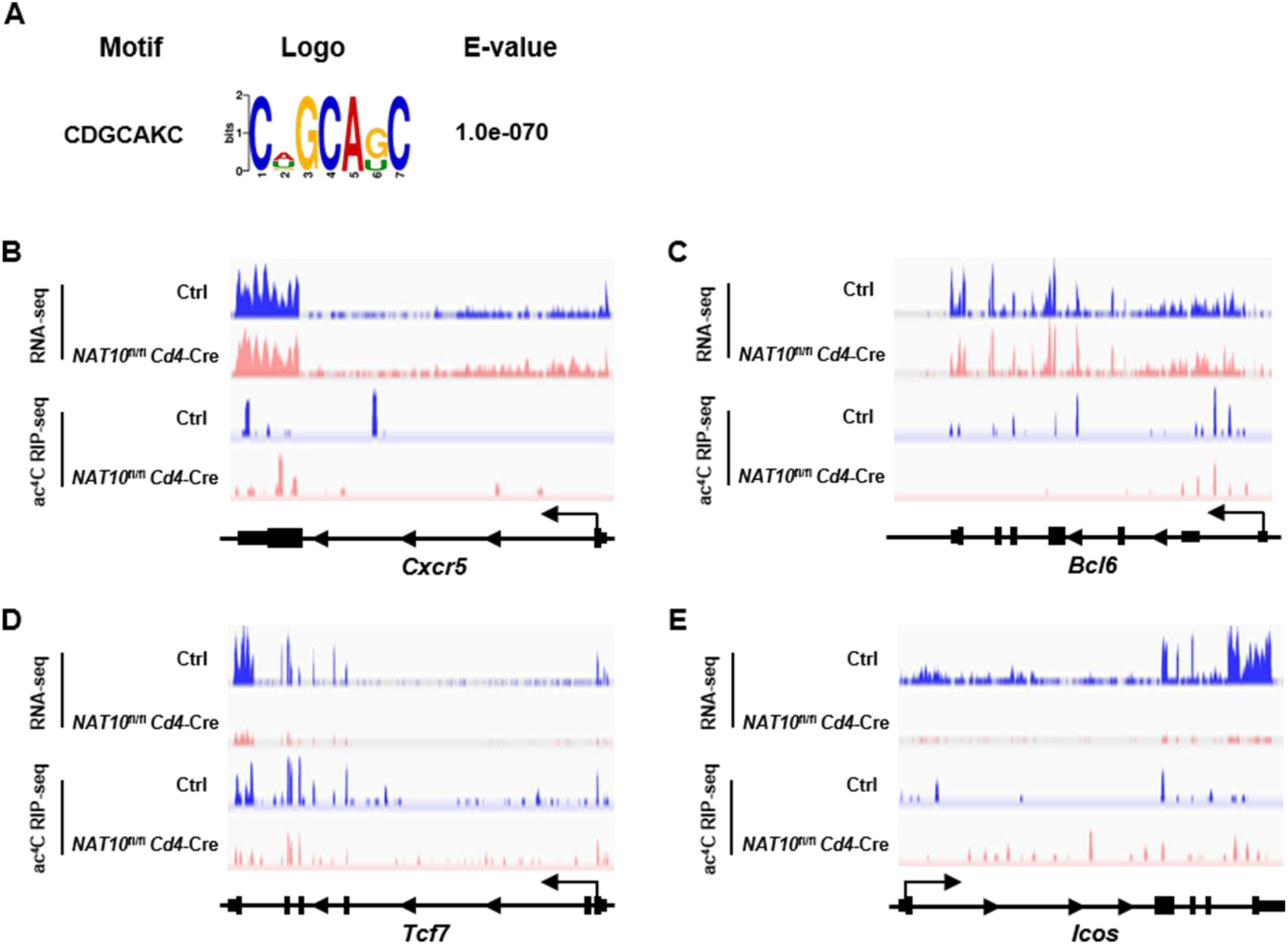
ac^4^C modification motif and ac^4^C distribution at genes. **A**, Putative ac^4^C acetylation motif. The enriched motifs and their E-values are shown. **B-E**, Integrative Genomics Viewer (IGV) tracks displaying RNA-seq (top panel) and ac^4^C RIP-seq (bottom panel) reads distribution of *Cxcr5*, *Bcl6*, *Tcf7* and *Icos* gene.

**Figure S7.**
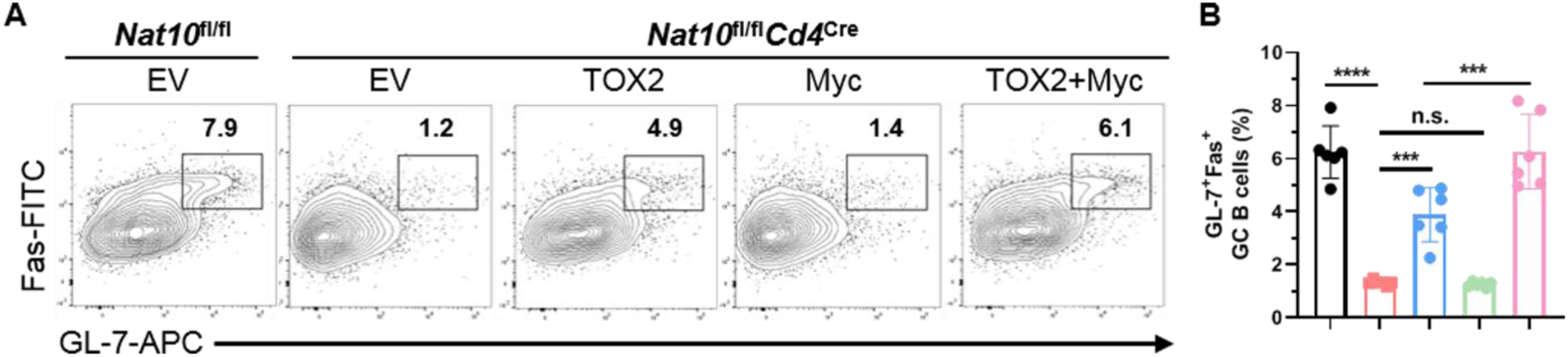
TOX2 overexpression rescues GC B cell responses in NAT10-deficient chimeras. **A-B**, Flow cytometry analysis of GL-7^+^Fas^+^ GC B cells gated on adoptive cells identified by CD45.2 staining in each group at 8 days post infection. Summary of the frequency and cell numbers of T_FH_ cells are shown in **B** (n = 6 per group). P values are calculated using two-tailed Student’s t-test for paired comparisons or one-way ANOVA for multiple comparisons. ***denotes p < 0.001 and ****denotes p < 0.0001 for the indicated comparison. n.s. = not significant.

**Figure S8.**
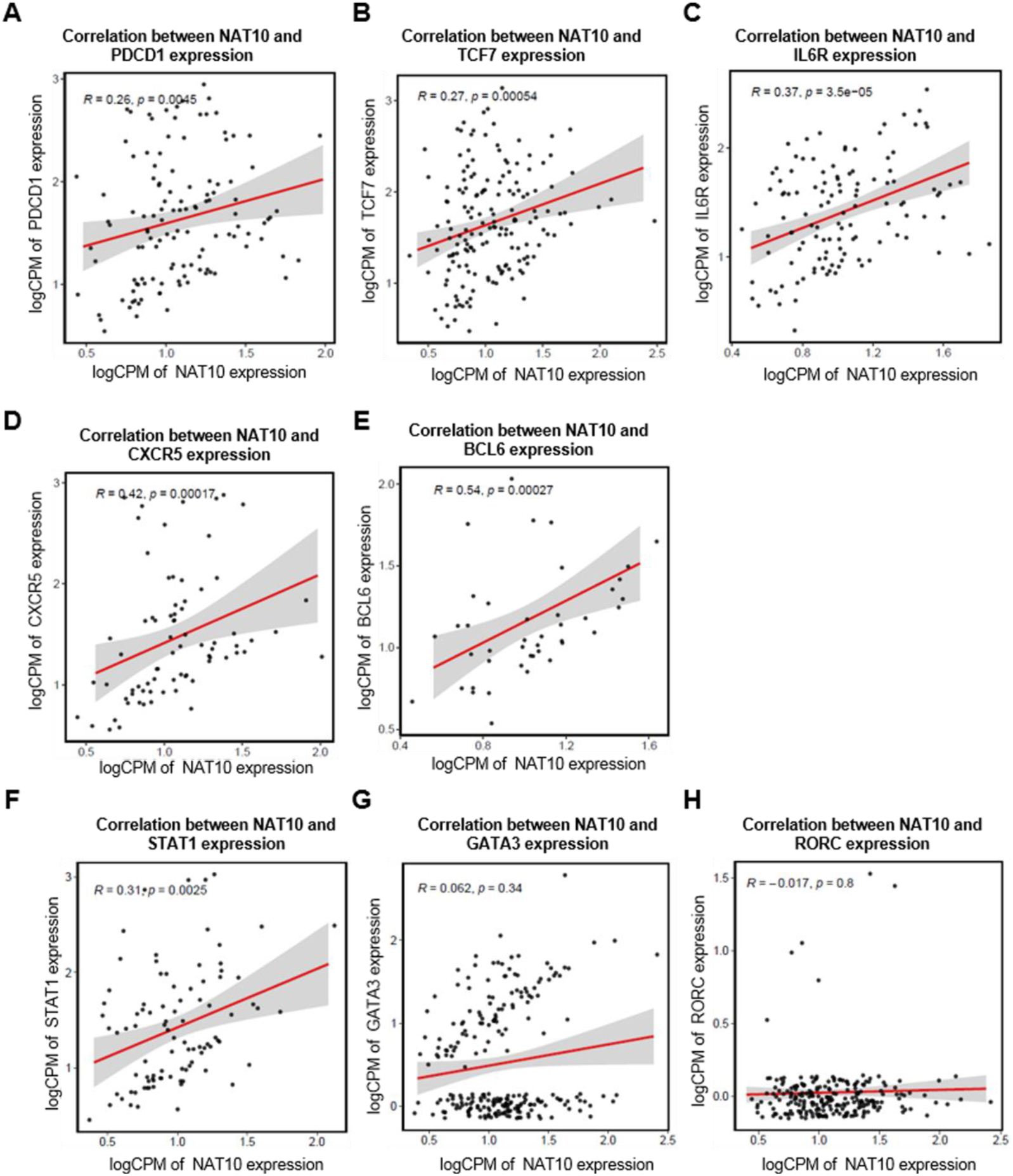
Correlation of NAT10 expression with T_FH_ signature in humans. **A-E**, Scatter plot comparing the mRNA expression of NAT10 and key T_FH_-associated genes in GC T_FH_ cells populations from human volunteers over one year period. **F-H**, Scatter plot comparing the NAT10 expression and T_H_1, T_H_2, T_H_17 marker genes (STAT1, GATA3 and RORC) from human volunteers over one year period. Correlation analyses were performed using the Pearson correlation metric, and P values were computed using two-sided t test.

**Figure S9.**
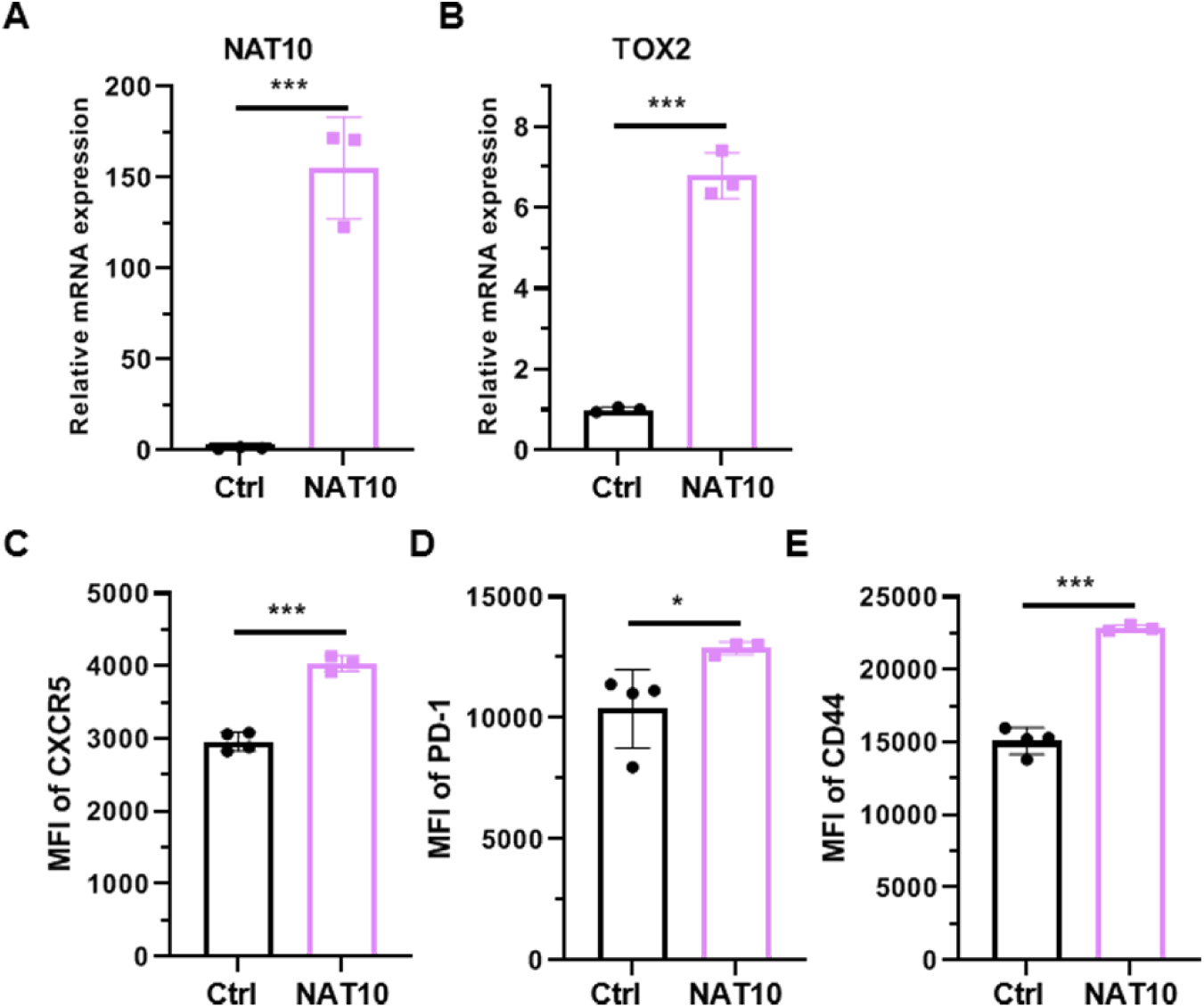
NAT10 overexpression enhances human T_FH_ cell differentiation in vitro. **A-B**, Human CD4^+^ T cells were activated *in vitro* under T_FH_ conditions and infected with Ctrl or NAT10 retrovirus, and the mRNA expression of NAT10 and TOX2 genes were determined by RT-qPCR. **C-E**, Flow cytometry analysis of MFI of CXCR5, PD-1 and CD44 on human T_FH_ cells activated *in vitro* under T_FH_ conditions (n = 3-4 per group). P values are calculated using two-tailed Student’s t-test for paired comparisons or one-way ANOVA for multiple comparisons. *denotes p < 0.05 and ***denotes p < 0.001 for the indicated comparison.

**Figure S10.**
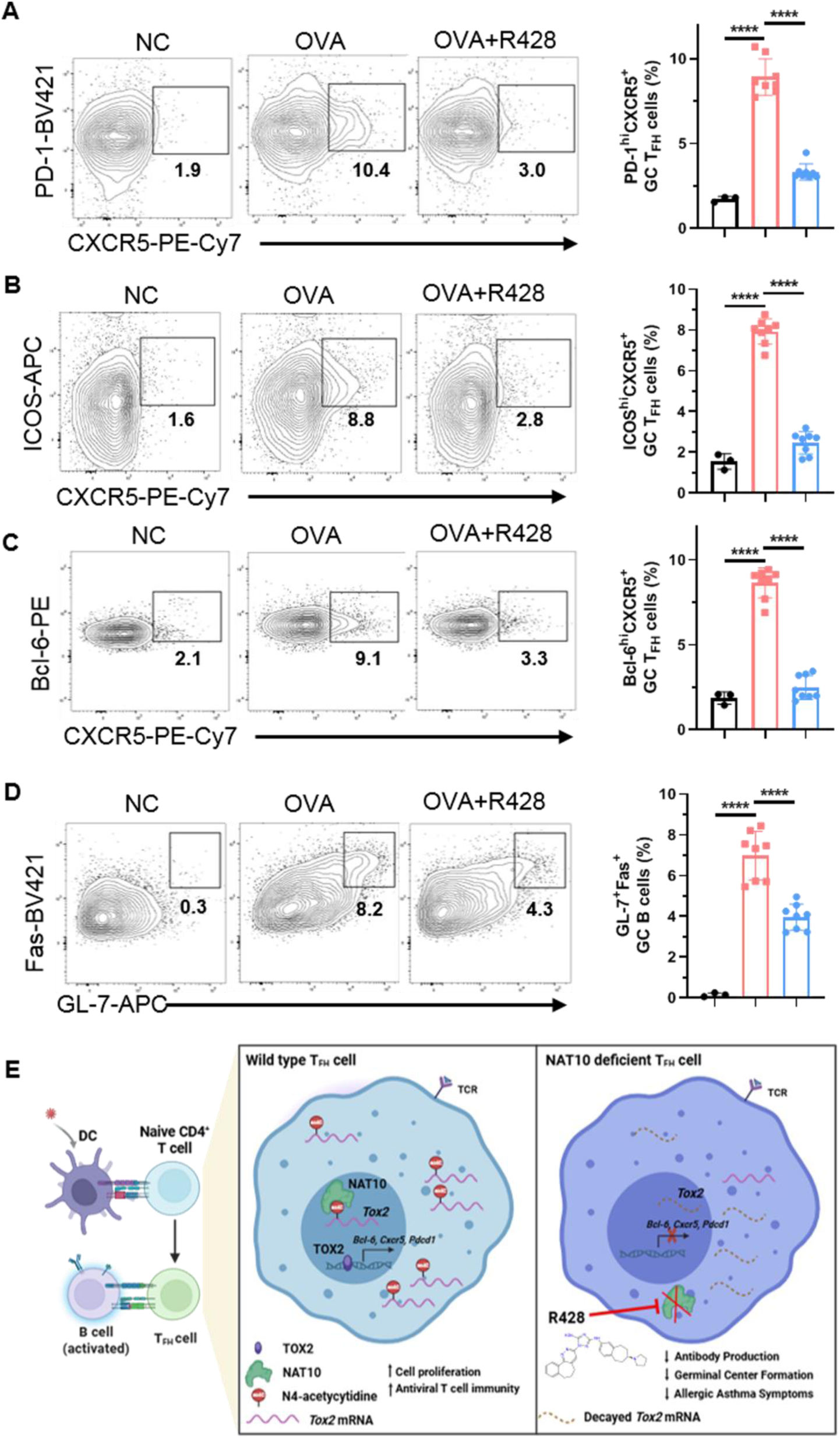
R428 treatment inhibits GC T_FH_ and B cell responses in asthma. **A**, Flow cytometry analysis and frequency of PD-1^hi^CXCR5^+^ GC T_FH_ cells, gated on CD4^+^ T cells, from mice with asthma treated with saline or R428 at day 24 (n =3-8). **B-C**, Flow cytometry analysis and frequency of ICOS^hi^CXCR5^+^ and Bcl-6^hi^CXCR5^+^ T_FH_ cells from from mice with asthma treated with saline or R428 at day 24 (n =3-8). **D**, Flow cytometry analysis of GL-7^+^Fas^+^ GC B cells from from mice with asthma treated with saline or R428 at day 24 (n =3-8). **E**, A proposed working model. P values are calculated using two-tailed Student’s t-test for paired comparisons or one-way ANOVA for multiple comparisons. ****denotes p < 0.0001 for the indicated comparison.

**Figure S11.**
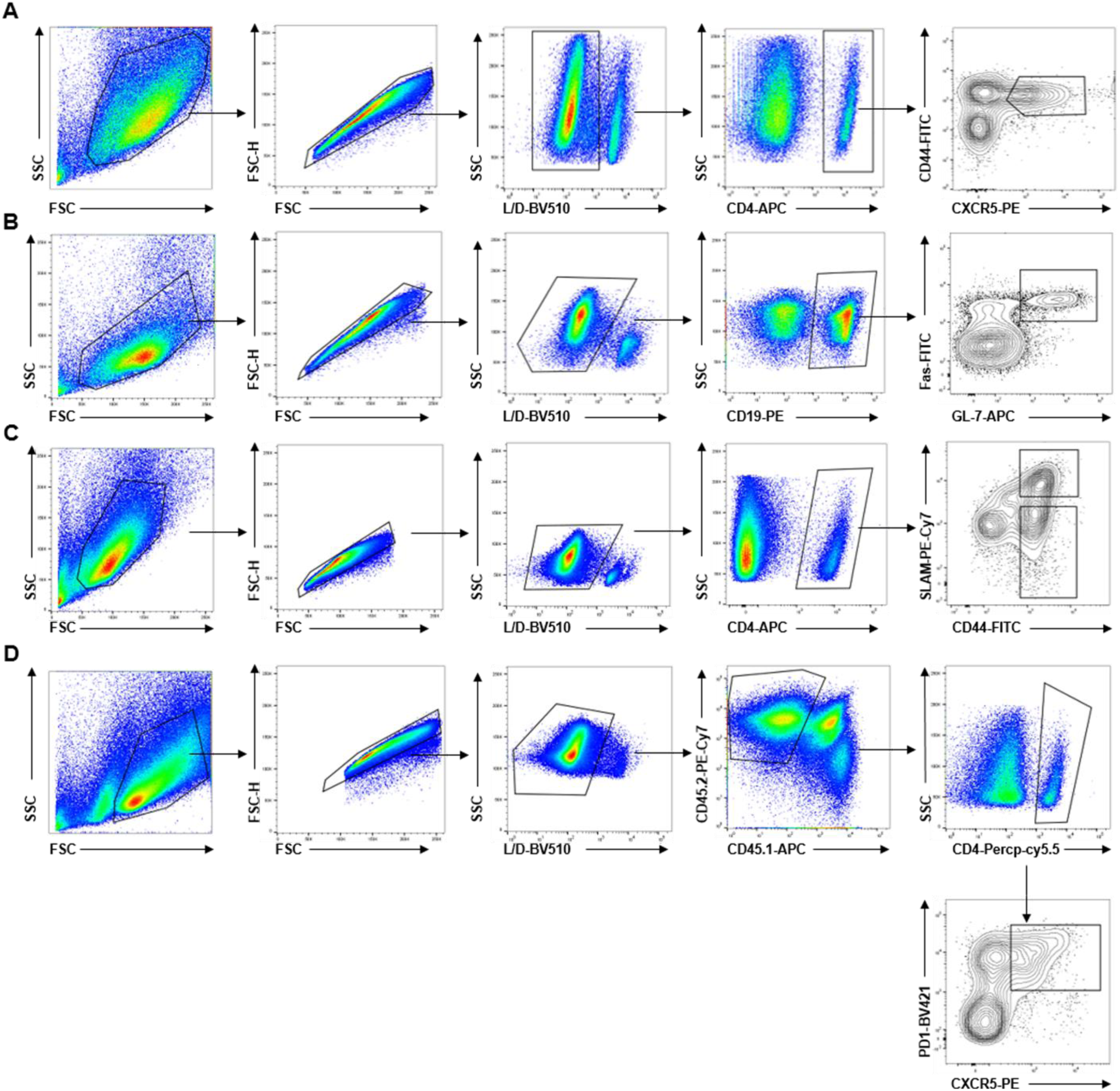
Gating strategy for flow cytometry analysis and cell sorting. **A**, Analysis of T_FH_ cell responses after LCMV-Armstrong infection (related to Figure 1 and S1). **B**, Analysis of GC B cells responses after LCMV-Armstrong infection (related to Figure 1 and S1). **C**, CD44^+^SLAM^hi^ T_H_1 cells and CD44^+^SLAM^lo^ T_FH_ cells were sorted for RNA-seq analysis (related to Figure 2 and S2). **D**, Representative flow cytometry gating strategies for T_FH_ cells deriving from transferred HSCs in chemic mice after LCMV-Armstrong infection (related to Figure 4 and S4).

## References

[1] Vinuesa C G, Linterman M A, Yu D, et al. Follicular Helper T Cells [J]. Annual review of immunology, 2016, 34: 335–68.

[2] Crotty S. Follicular helper CD4 T cells (Tfh) [J]. Annu Rev Immunol, 2011, 29: 621–63.

[3] Crotty S. T Follicular Helper Cell Biology: A Decade of Discovery and Diseases [J]. Immunity, 2019, 50(5): 1132–48.

[4] Vogelzang A, Mcguire H M, Yu D, et al. A fundamental role for interleukin-21 in the generation of T follicular helper cells [J]. Immunity, 2008, 29(1): 127–37.

[5] He J, Tsai L M, Leong Y A, et al. Circulating precursor CCR7(lo)Pd-1(hi) CXCR5(+) CD4(+) T cells indicate Tfh cell activity and promote antibody responses upon antigen reexposure [J]. Immunity, 2013, 39(4): 770–81.

[6] Walker L S K. The link between circulating follicular helper T cells and autoimmunity [J]. Nat Rev Immunol, 2022, 22(9): 567–75.

[7] Zander R, Kasmani M Y, Chen Y, et al. Tfh-cell-derived interleukin 21 sustains effector CD8(+) T cell responses during chronic viral infection [J]. Immunity, 2022, 55(3): 475–93 e5.

[8] Schmitt N, Bentebibel S E, Ueno H. Phenotype and functions of memory Tfh cells in human blood [J]. Trends Immunol, 2014, 35(9): 436–42.

[9] Morita R, Schmitt N, Bentebibel S E, et al. Human blood CXCR5(+)CD4(+) T cells are counterparts of T follicular cells and contain specific subsets that differentially support antibody secretion [J]. Immunity, 2011, 34(1): 108–21.

[10] Bentebibel S E, Lopez S, Obermoser G, et al. Induction of Icos+CXCR3+CXCR5+ Th cells correlates with antibody responses to influenza vaccination [J]. Sci Transl Med, 2013, 5(176): 176ra32.

[11] Xia Y, Sandor K, Pai J A, et al. BCL6-dependent Tcf-1(+) progenitor cells maintain effector and helper CD4(+) T cell responses to persistent antigen [J]. Immunity, 2022, 55(7): 1200–15 e6.

[12] Choi Y S, Gullicksrud J A, Xing S, et al. Lef-1 and Tcf-1 orchestrate T(Fh) differentiation by regulating differentiation circuits upstream of the transcriptional repressor Bcl6 [J]. Nat Immunol, 2015, 16(9): 980–90.

[13] Xu L, Cao Y, Xie Z, et al. The transcription factor Tcf-1 initiates the differentiation of T(Fh) cells during acute viral infection [J]. Nat Immunol, 2015, 16(9): 991–9.

[14] Ise W, Kohyama M, Schraml B U, et al. The transcription factor Batf controls the global regulators of class-switch recombination in both B cells and T cells [J]. Nat Immunol, 2011, 12(6): 536–43.

[15] Johnston R J, Poholek A C, Ditoro D, et al. Bcl6 and Blimp-1 are reciprocal and antagonistic regulators of T follicular helper cell differentiation [J]. Science, 2009, 325(5943): 1006–10.

[16] Guo G, Shi X, Wang H, et al. Epitranscriptomic N4-Acetylcytidine Profiling in CD4(+) T Cells of Systemic Lupus Erythematosus [J]. Front Cell Dev Biol, 2020, 8: 842.

[17] Yu C, Chen Y, Luo H, et al. NAT10 promotes vascular remodelling via mrna ac4c acetylation [J]. Eur Heart J, 2025, 46(3): 288–304.

[18] Sun L, Li X, Xu F, et al. A critical role of N(4)-acetylation of cytidine in mrna by NAT10 in T cell expansion and antiviral immunity [J]. Nat Immunol, 2025.

[19] Arango D, Sturgill D, Alhusaini N, et al. Acetylation of Cytidine in mrna Promotes Translation Efficiency [J]. Cell, 2018, 175(7): 1872–86 e24.

[20] Harada Y, Tanaka S, Motomura Y, et al. The 3’ enhancer CNS2 is a critical regulator of interleukin-4-mediated humoral immunity in follicular helper T cells [J]. Immunity, 2012, 36(2): 188–200.

[21] Kobayashi T, Iijima K, Dent A L, et al. Follicular helper T cells mediate Ige antibody response to airborne allergens [J]. J Allergy Clin Immunol, 2017, 139(1): 300–13 e7.

[22] Sun L, Li X, Xu F, et al. A critical role of N(4)-acetylation of cytidine in mrna by NAT10 in T cell expansion and antiviral immunity [J]. Nat Immunol, 2025, 26(4): 619–34.

[23] Fahey L M, Wilson E B, Elsaesser H, et al. Viral persistence redirects CD4 T cell differentiation toward T follicular helper cells [J]. J Exp Med, 2011, 208(5): 987–99.

[24] Harker J A, Lewis G M, Mack L, et al. Late interleukin-6 escalates T follicular helper cell responses and controls a chronic viral infection [J]. Science, 2011, 334(6057): 825–9.

[25] Angum F, Khan T, Kaler J, et al. The Prevalence of Autoimmune Disorders in Women: A Narrative Review [J]. Cureus, 2020, 12(5): e8094.

[26] Li H, Cai X, Xu C, et al. Rna cytidine acetyltransferase NAT10 maintains T cell pathogenicity in inflammatory bowel disease [J]. Cell Discov, 2025, 11(1): 19.

[27] Wang J N, Suo X G, Yu J T, et al. NAT10 exacerbates acute renal inflammation by enhancing N4-acetylcytidine modification of the CCL2/CXCL1 axis [J]. Proc Natl Acad Sci U S A, 2025, 122(17): e2418409122.

[28] Wu T, Shin H M, Moseman E A, et al. TCF1 Is Required for the T Follicular Helper Cell Response to Viral Infection [J]. Cell Rep, 2015, 12(12): 2099–110.

[29] Tibbitt C A, Stark J M, Martens L, et al. Single-Cell Rna Sequencing of the T Helper Cell Response to House Dust Mites Defines a Distinct Gene Expression Signature in Airway Th2 Cells [J]. Immunity, 2019, 51(1): 169–84 e5.

[30] Ciofani M, Madar A, Galan C, et al. A validated regulatory network for Th17 cell specification [J]. Cell, 2012, 151(2): 289–303.

[31] Liu X, Yan X, Zhong B, et al. Bcl6 expression specifies the T follicular helper cell program in vivo [J]. J Exp Med, 2012, 209(10): 1841–52, S1-24.

[32] Xu W, Zhao X, Wang X, et al. The Transcription Factor Tox2 Drives T Follicular Helper Cell Development via Regulating Chromatin Accessibility [J]. Immunity, 2019, 51(5): 826–39 e5.

[33] Schattgen S A, Turner J S, Ghonim M A, et al. Influenza vaccination stimulates maturation of the human T follicular helper cell response [J]. Nat Immunol, 2024, 25(9): 1742–53.

[34] Xu Z, Zhu M, Geng L, et al. Targeting NAT10 attenuates homologous recombination via destabilizing Dna:Rna hybrids and overcomes Parp inhibitor resistance in cancers [J]. Drug Resist Updat, 2025, 81: 101241.

[35] Larrieu D, Britton S, Demir M, et al. Chemical inhibition of NAT10 corrects defects of laminopathic cells [J]. Science, 2014, 344(6183): 527–32.

[36] Zhang S, Huang F, Wang Y, et al. NAT10-mediated mrna N(4)-acetylcytidine reprograms serine metabolism to drive leukaemogenesis and stemness in acute myeloid leukaemia [J]. Nat Cell Biol, 2024, 26(12): 2168–82.

[37] Horiuchi S, Wu H, Liu W C, et al. Tox2 is required for the maintenance of Gc T(Fh) cells and the generation of memory T(Fh) cells [J]. Sci Adv, 2021, 7(41): eabj1249.

[38] Holland S J, Pan A, Franci C, et al. R428, a selective small molecule inhibitor of Axl kinase, blocks tumor spread and prolongs survival in models of metastatic breast cancer [J]. Cancer Res, 2010, 70(4): 1544–54.

[39] Dangi T, Chung Y R, Palacio N, et al. Interrogating Adaptive Immunity Using Lcmv [J]. Curr Protoc Immunol, 2020, 130(1): e99.

[40] Ahmed R, Salmi A, Butler L D, et al. Selection of genetic variants of lymphocytic choriomeningitis virus in spleens of persistently infected mice. Role in suppression of cytotoxic T lymphocyte response and viral persistence [J]. J Exp Med, 1984, 160(2): 521–40.

[41] Wolock S L, Lopez R, Klein A M. Scrublet: Computational Identification of Cell Doublets in Single-Cell Transcriptomic Data [J]. Cell Syst, 2019, 8(4): 281–91 e9.

[42] Wolf F A, Angerer P, Theis F J. Scanpy: large-scale single-cell gene expression data analysis [J]. Genome Biol, 2018, 19(1): 15.

[43] Dai M, Pei X, Wang X J. Accurate and fast cell marker gene identification with Cosg [J]. Brief Bioinform, 2022, 23(2).

[44] Cho K S, Park M K, Kang S A, et al. Adipose-derived stem cells ameliorate allergic airway inflammation by inducing regulatory T cells in a mouse model of asthma [J]. Mediators Inflamm, 2014, 2014: 436476.

[45] Wu J, Wang P, Xie X, et al. Gasdermin D silencing alleviates airway inflammation and remodeling in an ovalbumin-induced asthmatic mouse model [J]. Cell Death Dis, 2024, 15(6): 400.

[46] De Castro L L, Xisto D G, Kitoko J Z, et al. Human adipose tissue mesenchymal stromal cells and their extracellular vesicles act differentially on lung mechanics and inflammation in experimental allergic asthma [J]. Stem Cell Res Ther, 2017, 8(1): 151.

[47] Padrid P, Snook S, Finucane T, et al. Persistent airway hyperresponsiveness and histologic alterations after chronic antigen challenge in cats [J]. Am J Respir Crit Care Med, 1995, 151(1): 184–93.

